# The Cause of Most Common Antler Abnormality is Not a Fracture But an Osteomyelitis Like Disease the Pedunculitis Chronica Deformans

**DOI:** 10.1101/2025.02.05.636259

**Authors:** Farkas Sukosd, Istvan Lakatos, Adam Urmos, Bianka Babarczi, Gabriella Skoda, Zsofia Molnar, Tibor Pankotai, Barbara N Borsos, Reka Karkas, Akos Sukosd, Gabor Palanki, Szabolcs Dobrosy, Attila Arany Toth, Karoly Erdelyi, Gyozo Horvath, Adrienn Horvath, Daniel Toth, Mate Maurer, Mihaly Miso, Patrik Plank, Arpad Czeh, Gyorgy Nagyeri, Szilamer Ferenczi, Peter Gobolos, Gabor Kemenesi, Katalin Posta, Ferenc Kovacs, Miklos Mezes, Laszlo Szemethy, Zsuzsanna Szoke

## Abstract

Climate change is increasing the incidence of certain diseases in wildlife, offering a deeper understanding of their underlying causes. Traditionally, the most common antler abnormalities have been attributed to trauma, based on their morphology. However, this explanation does not account for the epidemic-like spread of abnormalities in specific regions, suggesting that they might be a previously undescribed aspect of this anomaly. In fallow deer (*Dama dama*), using advanced medical diagnostic tools, we identified an osteomyelitis-like condition, Pedunculitis Chronica Deformans (PCD) which is also observed in other *Cervidae*. The incidence of this disease shows a strong correlation with the production of mycotoxins by molds that are becoming more prevalent with global warming. Mycotoxins can inhibit wound healing through scar formation after the antlers are cast, leading to the development of PCD. In this report we characterize PCD pathomorphology and introduce a scoring system to differentiate it from other deformities. Moreover, we argue that the most common anomalies such as fractures, antler losses, fatal meningoencephalitis and brain abscesses are complications of PCD. Our research underscores the potential of antlers to serve as biomarkers for mycotoxin exposure.

## Introduction

Antlers are unique outgrowths specific to the *Cervidae* family that reflect the biological condition of the animal, exhibiting inherited phenotypes, nutritional status, environmental impacts and the quality of wildlife management ^1,2^. Notably, antlers represent the fastest-growing and regenerating bone tissue found in nature^1^. While antler abnormalities are rare ^3^, they hold significant value as trophies among hunters. The earliest scientific publication on antler deformities dates back to 1900^4^.

Hungary is recognized internationally for the high quality of its deer population, with two world-record fallow deer (*Dama dama*) trophies originating from its Northeast Region^5^. However, since the latter half of the 20th century a notable increase in antler deformities has been observed in the Southern regions of the country, where they are now considered endemic. In the affected areas, 40-80% of harvested male fallow deer^6^, alongside increasing numbers of red deer (*Cervus elaphus)* and roe deer (*Capreolus capreolus*) display some form of abnormality. Additionally, a rising incidence of such abnormalities has been noted among the elk (*Cervus canadensis*) population in Northern Arizona, including the Hualapai Indian Reservation^7^. Instances of comparable malformations were also documented on mule deer (*Odocoileus hemionus*) in central Utah in the mid-20th century,^8^ as well as in North-Central Kansas, where extreme antler malformations identical in morphology have been noted^9^. Particularly concerning are the distinctive osseous deformities involving the pedicle - the frontal bone outgrowth serving as the basis of the antler- and adjacent skull region. Such anomalies have been documented in white-tailed deer (*Odocoileus virginianus*) from Georgia, US where they were linked to peri-peduncular purulent inflammation and associated meningoencephalitis or intracranial abscess complications ^10,11^. A comprehensive study conducted across 12 US states and four Canadian provinces found that 2.2% of 4,500 white-tailed deer exhibited central nervous system inflammation coupled with pedicle, skull, and antler abnormalities^10^. This highly spatially variable disease was detected in 9 (35%) cases in a radio-collared male white-tailed deer population of 26 individuals aged 2.5 years or older in Kent County, Maryland, USA^12^. Despite these findings, the true extent and epidemiology of antler deformities remains unclear. Without precise descriptions, it is not possible to distinguish between pathological abnormalities and physiological variants. In addition, trophy evaluation procedures exclude cases showing such deformities, as the specimens concerned do not meet the criteria for standard evaluation. Consequently, the extent of these conditions remains under-reported and poorly understood.

Mycotoxins (MTs) are persistent natural organic pollutants categorized as secondary metabolic products of plant-pathogenic fungi and are known to have a wide range of detrimental biological effects ^13,14^. However, their potential role in severe antler disorders has not yet been investigated. Aflatoxins (AFs) - produced by *Aspergillus flavus* and *A. parasiticus* -, are both genotoxic and immunotoxic, causing tissue damage by disrupting cell cycle processes^15^. Zearalenone (ZEA), a phytoestrogenic toxin from *Fusarium* species binds directly to estrogen receptors^16^, mimicking hormonal activity and potentially affecting reproductive and developmental processes. Deoxynivalenol (DON), the most prevalent *Fusarium* mycotoxin, is known to induce both acute and chronic mycotoxicosis, with dividing cells being particularly vulnerable compared to more differentiated cells^17^. Furthermore, fumonisins - another group of toxins from *Fusarium* - are known to induce apoptosis in liver cells, with the toxic effects significantly amplified when combined with AFs exposure^18^.

The level of mycotoxin (MT) contamination in a plant is highly influenced by various physicochemical factors, including environmental temperature, moisture, relative air humidity, and oxygen levels^19^. Studies have demonstrated that increased temperatures, elevated CO₂ levels, and other factors associated with climate change affect the frequency, distribution and accumulation of mycotoxin production. This, in turn, heightens the exposure risk for organisms, amplifying the potential impact of these toxins ^20,21^.

To investigate the rising incidence of antler anomalies in Hungarian *Cervidae* populations - including *Dama dama* (DD), *Capreolus capreolus* (CC) and *Cervus elaphus* (CE) - we described a chronic inflammatory disease - named *Pedunculitis Chronica Deformans* (PCD) - of the pedicle which is strongly associated with MT exposure. In this study we introduce a morphological scoring system for this chronic osteomyelitis-like disease providing a reliable method for differentiating it from other antler abnormalities caused by combat trauma, hormonal insufficiency, or injuries sustained during the velvet phase. Our work includes a detailed characterization of the *Peduncular-Dermal Junction* (PDJ) as a distinct anatomical structure, highlighting its significance as a key factor in the disease’s initiation and progression. Further, we explored the hormonal influences and the role of MTs in the pathomechanism of PCD, detailing how these factors can bring about complications such as bacterial superinfections, which can escalate into life-threatening conditions like meningoencephalitis and brain abscesses. Our research sheds new light on a long-misunderstood antler anomaly, revealing that its prevalence has accelerated significantly in recent decades. This study provides a more comprehensive understanding of the underlying causes of these deformities and establishes a foundation for future diagnostics and treatment.

## Results and discussion

### Epidemiology, age distribution and healthy status of Cervidae with aberrant trophies

Over the past decade, the rate of antler abnormalities in Hungary (Central Europe) has increased dramatically, to a degrees that is striking even to the casual observer **(Fig. 1, Supplementary Fig. 1 and Supplementary Video 1)**. The analysis of our Southern Hungarian non species-selected aberrant trophy collection in the period from 2017 to 2021 showed significant increase in antler anomalies in various free living Cervidae species. The incidence of aberrant trophies among fallow deer (*Dama dama*, DD) was notably higher than in roe deer (*Capreolus capreolus*, CC) and red deer (*Cervus elaphus*, CE), with ten times more anomalies in DD compared to CC (50 vs. 5) and approximately 16 times more than CE (50 vs. 3). When adjusted for population density, DD appears to be impacted the most by these deformities: 34 times more frequently than CC and 50 times more than CE, suggesting a unique vulnerability. **(Supplementary Data 1).** Therefore, our detailed analysis specifically focused on DD, the data related to alterations in CC **(Supplementary Section 1, Supplementary Fig 2 and SupplementaryData 2)** and CE **(Supplementary Section 2, Supplementary Fig 3 and SupplementaryData 3)** can be found in the Supplement.

**Fig. 1.**
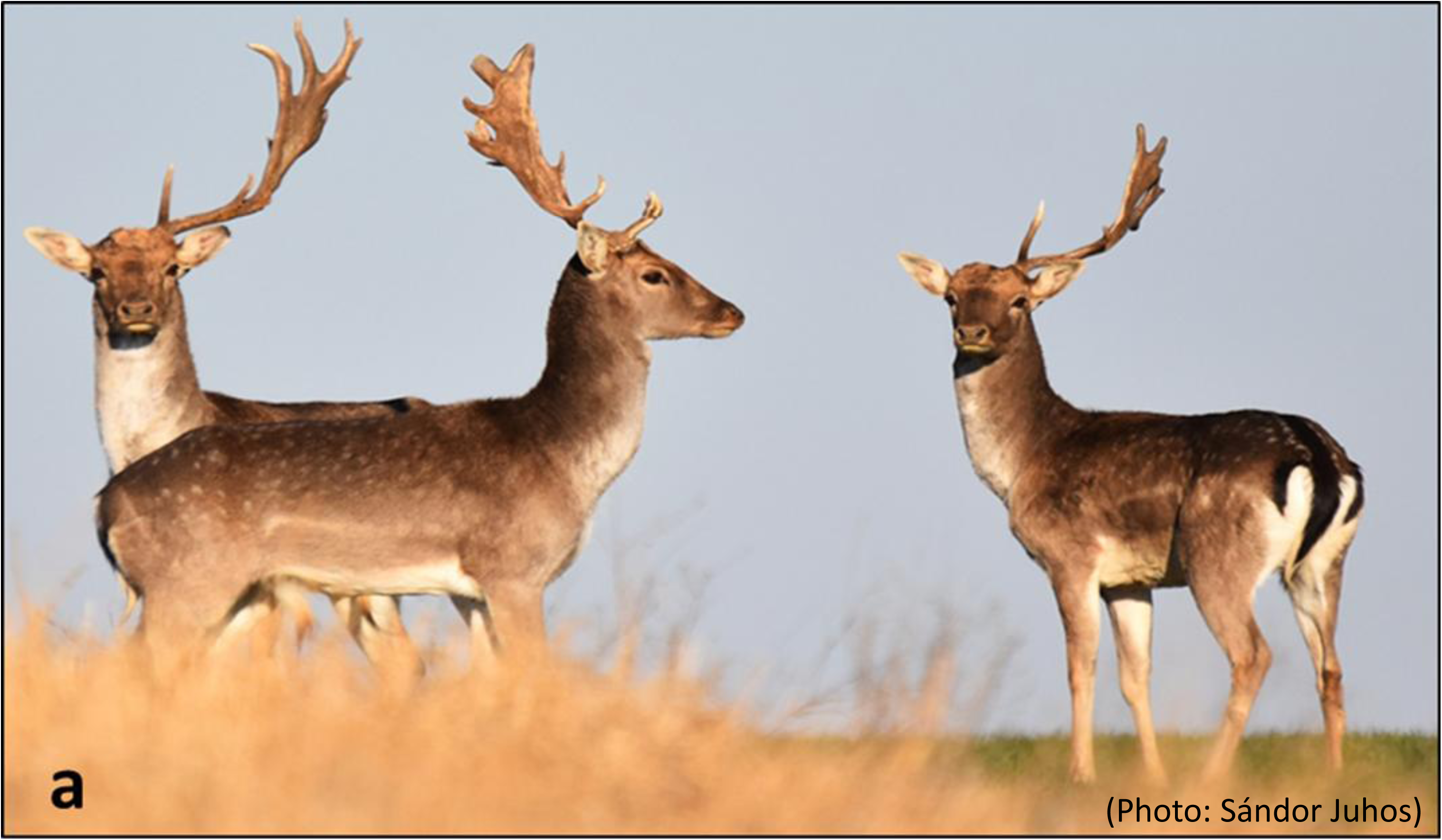
Unfortunately, it is not a rare picture from the area of Kocsola (Southern Hungary, Central Europe) showing simultaneously three fallow deer (*Dama dama*) with antler problems. They keep their heads straight, which shows that the loss of antlers did not happen during the fight, but that they could not develop them. They had to compensate for this weight asymmetry by neck muscle hypertrophy for the purpose of reaching the normal head position and vision which take a long time. (Photo: Sándor Juhos). More nature photos with the same phenomena can be found in the supplement.

In the Hungarian Trophy Register (HTR), data on age, main beam and brow tine sizes are recorded for normal trophies **(Fig. 2F)** from 2017 to 2020, a total of 2,924 register sheets documented these parameters regarding our study region. Of these, 12 cases (0.4%) included pedicle abnormalities and were excluded from analysisThe remaining 2912 records formed our Trophy Register Control Group for brow tine and main beam sizes across age groups. This low incidence rate in records contrasts sharply with the 40-80% frequency of abnormalities observed during harvests in this region^6^ **(Fig. 2A)**. The discrepancy may be due to the lack of standardized methods for describing malformed trophies. Scoring systems such as those by the International Council for Game and Wildlife Conservation (CIC score, https://www.cic-wildlife.org) and Boone and Crockett Club (B&C score https://www.boone-crockett.org) are well-suited for assessing healthy trophy sizes and forms but lack guidelines for classifying malformations (**Fig. 2B**). Consequently, only sporadic notes on these anomalies appear in trophy evaluations. The remaining 2,900 healthy cases formed our Trophy Register Control Group (TRCG), with an average age of 9 years, compared to an average age of 7.74 years for aberrant DD trophies (p<0.0001). Kaplan-Meier survival analysis revealed reduced life expectancy for animals with deformed antlers (**Fig. 2C**). For the analysis of health, toxicological and hormonal status, samples were collected from 58 abnormal antlered animals and 31 control animals during the same hunting season and their condition was documented in the hunting records. Animals with deformed antlers showed significantly poorer condition as per Hungarian Body Condition Score evaluation criteria (Fisher’s test, p<6,51*10^-6^) (**Supplementary Data 4**). Although hunting-related mortality is influenced by planned culling, it is still associated with the animal’s health status. Our findings are consistent with the observations that animals with antler deformities are often culled earlier due to poor health rather than the value of their trophies ^22,23^. Evidence indicates that individuals with deformed antlers across all age groups have reduced life expectancy compared to those with normal antlers (**Fig. 2D and 2E**). Therefore, antler deformity should be identified as a sign of serious disease.

**Fig. 2.**
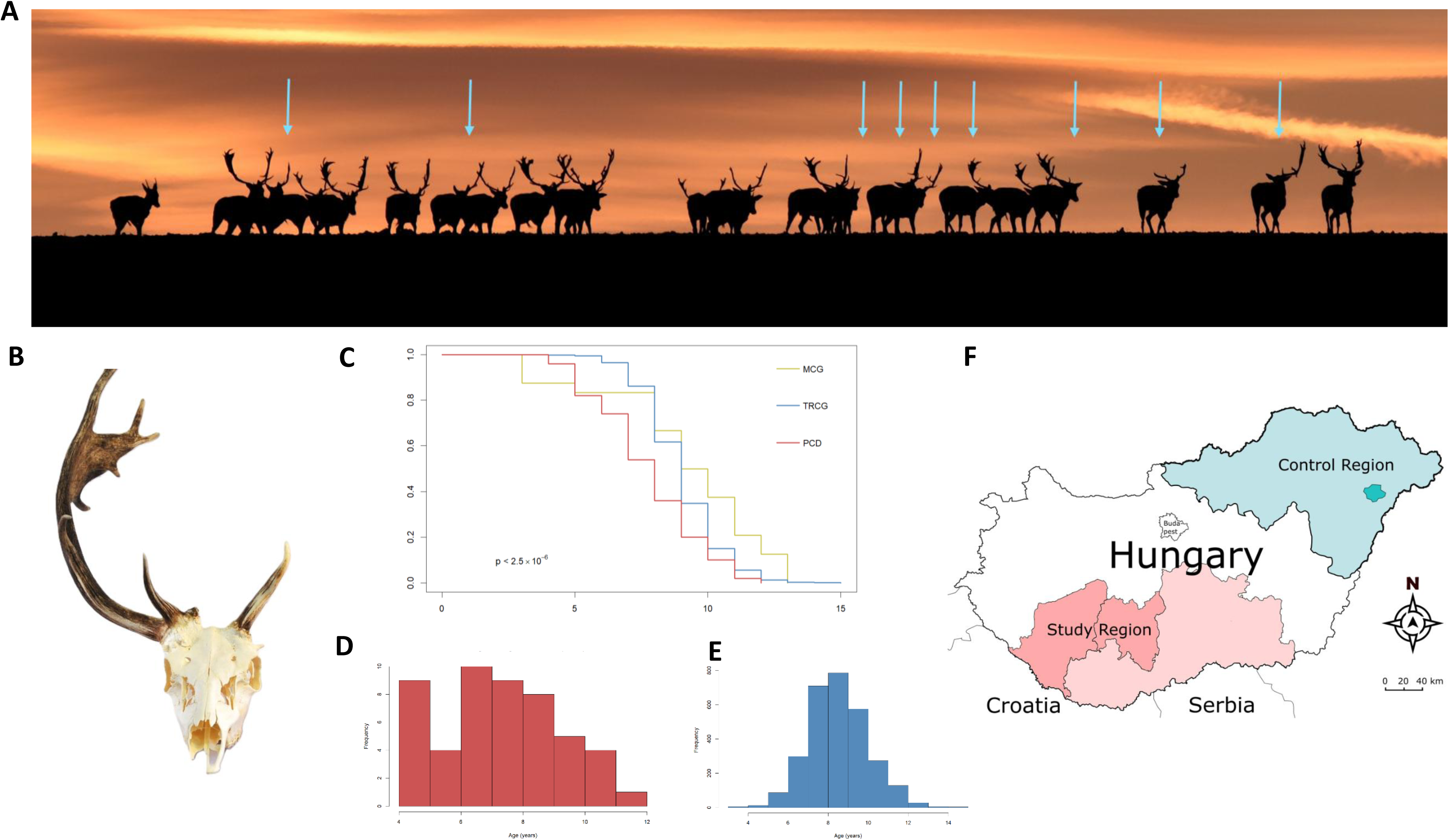
Antler abnormality in Southern Hungary. **A**. An herd of fallow deer *(Dama dama,* DD) at sunset, from the area of Kocsola (Southern Hungary) (Photo: Sándor Juhos). Nine out of 23 (39%) show antler abnormalities visible from a distance. **B.** A typical example of aberrant DD trophy. These cases are excluded from the classical trophy evaluation, so there are no reliable data on their proportion**. C.** Although hunting-related mortality is influenced by planned culling, it is still associated with the animal’s health status. Kaplan-Meier survival analysis revealed reduced life expectancy for animals with deformed antlers compeared with healthy member of Morphological (MCG) and Trophy Register (TRCG) Controll Group. **D** Age distribution of the diseased individuals, with an average age of 7.7 years. **E** Compared to the age distribution of the TRCG, where the average age was 9 years. (p<0.0001). **F** The test samples came from the two southern regions of Hungary (Southern Transdanubia and Southern Great Plain) and the controls from the north-eastern region (Northern Hungary). The areas of Croatia and Serbia related to the study region are also indicated.

### Peduncular-Dermal Junction (PDJ); Structural and Pathological Insights

In diseased individuals, we observed a distinct macroscopic separation between the peripeduncular soft tissues—including the skin, subcutaneous connective tissue, and periosteum—and the underlying bone (**Fig. 3h**). This separation suggests that damage to a critical peripheral structural tight junction around the pedicle may contribute significantly to disease development.

**Fig. 3.**
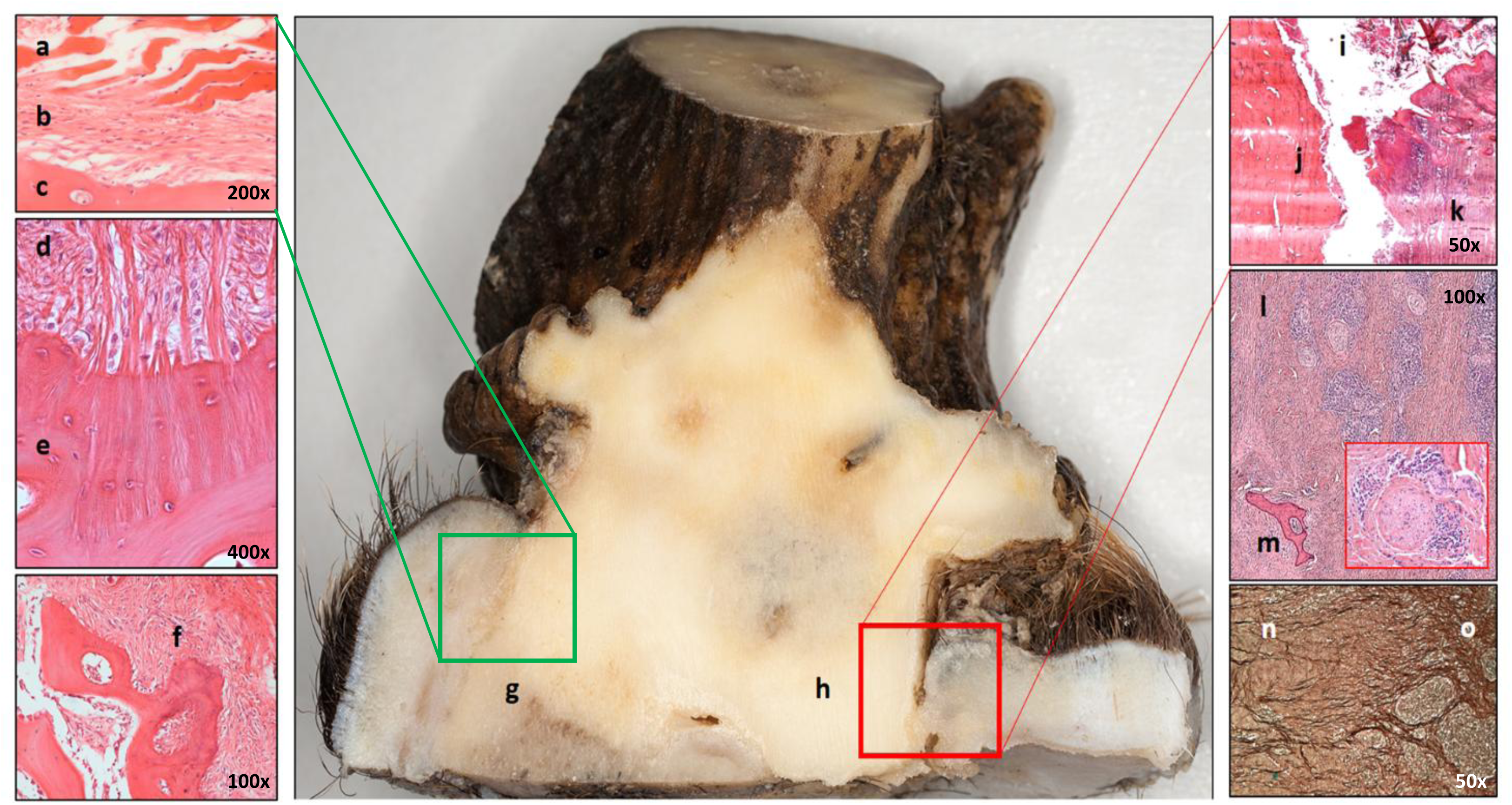
The Peduncular Dermal Junction (PDJ) and its damage on saggitaly sectioned peduncle and peripeduncular soft tissue. Green squares indicate healthy and red, diseased areas from which excisions were made for microscopic examination. **a-g** On the left, the histological components of the PDJ are shown. The epidermis (a) is directly attached to the bone (c) surface without dermis (b). **d-e** A Sharpey fibres anchor the periosteum (d) to the bone (e)**. f** At lower magnification, the bone surface roughness is visible, which increases the adhesion surface. The trabecular bone is on the left side of the picture, the periosteum on the right. **h-o** Damage to the PDJ caused periosteal bone discohesion (h), allowing the accumulation of foreign, contaminated substances, such as plant fibres, purulent exudate and tissue debris, in the resulting gap (i). Signs of superficial osteomyelitis are a surface covered with devitalized bone particles (j) and an accumulation of inflammatory cells in the skin, visible as a bluish discoloration (k). In the deeper soft tissue, a massive, predominantly perineural (see insert) and perivascular arrangement of lymphoplasmacytic inflammatory cells infiltration (l) with bone sequestrum (m) is seen. Chronic inflammation is characterized by fibrosis (n), which causes damage to the elastic fiber matrix (o) and disruption of Sharpey’s fibers, resulting in further enlargement of the gap with shrinkage of the peripeduncular tissue.

In mammals, the only hitherto characterised bony structure that penetrates the integument is the tooth, whose anatomical structure, especially at the periodontal junction is well-discussed^24^. This junction isolates the heterogeneous microbial flora of the oral cavity from the rest of the body, through the specialized structure of the oral mucosa^24^. Antlers in *Cervidae* are bony structures that penetrate the integument when the velvet is shed and the continuity of the skin is interrupted^25^. This breach in the integument—the body’s first line of defense—potentially exposes the animal to pathogens from the surrounding environment, particularly around the dry antler. We described the soft tissue components around the pedicle as an independent structure referred to as the pedunculo-dermal junction (PDJ), which we believe plays a crucial role in maintaining integumentary integrity. Microscopically, the PDJ is composed of three primary elements: (1) The direct connection of the distal edge of the epidermis without an intervening dermis or subcutaneous layer to the pedicle periosteum ^26^; (2) Sharpey’s fibers anchoring the periosteum firmly to the bone ^27^ and (3) the microscopic roughness of the distal peduncular bone surface which is augmenting the contact area and potentially enhancing barrier strength **(Fig. 3 a-g)**. Li et al. demonstrated that the distal third of the pedicle periosteum is more resistant to detachment than the proximal two-thirds^28^ supporting our hypothesis that the PDJ is an independent histomorphological structural unit. In other words, segments of peripeduncular soft tissue of different heights have different significance not only in antler regeneration, ^29^ but also in the closure of the integument after casting of the velvet.

We found that damage to the PDJ caused periosteal-bone discohesion, allowing the accumulation of external contaminants, such as plant fibers in the resulting gap and introducing pathogens which bring about superinfections triggering acute inflammation characterized by purulent exudate and tissue debris. The fibrotic remodelling of connecting tissue leads to shrinkage of the surrounding elastic fiber matrix and the disruption of Sharpey’s fibers creating a favorable environment to chronic inflammation around the pedicle **(Fig. 3 h-o)**. The process can continue for several years, as evidenced by the case we identified in the velvet phase (**Supplementary Fig. 4**). The pathomechanism bears similarities to parodontitis, where persistent inflammation is exacerbated by compromised structural integrity ^24^.

### Characterization of the Aberrant Trophies

Aberrant trophies exhibit pathological variations not observed under physiological conditions, for example significant absence of rose granularity, remarkable differences between beam lengths, pedicle deformities, and skull abnormalities. Additionally, they often show the accumulation of unique anatomical variations rarely observed under normal circumstances, such as supernumerary tines. To characterize these abnormalities, which we dubbed RAPS features, we examined three malformations of the antler rose (R1-R3), six of the antler (A1, A2, A2.1, A3, A3.1, A4), seven of the pedicle (P1-P7), and six of the surrounding skull (S1-S6), altogether 22 characteristics. Since specific combinations of RAPS symptoms can also indicate disease severity, we established four grades of disease progression **(Fig. 4 and Supplementary Data. 5)** based on this framework which can serve as a starting point for a consensus classification based on a larger cohort in the future.

**Fig. 4.**
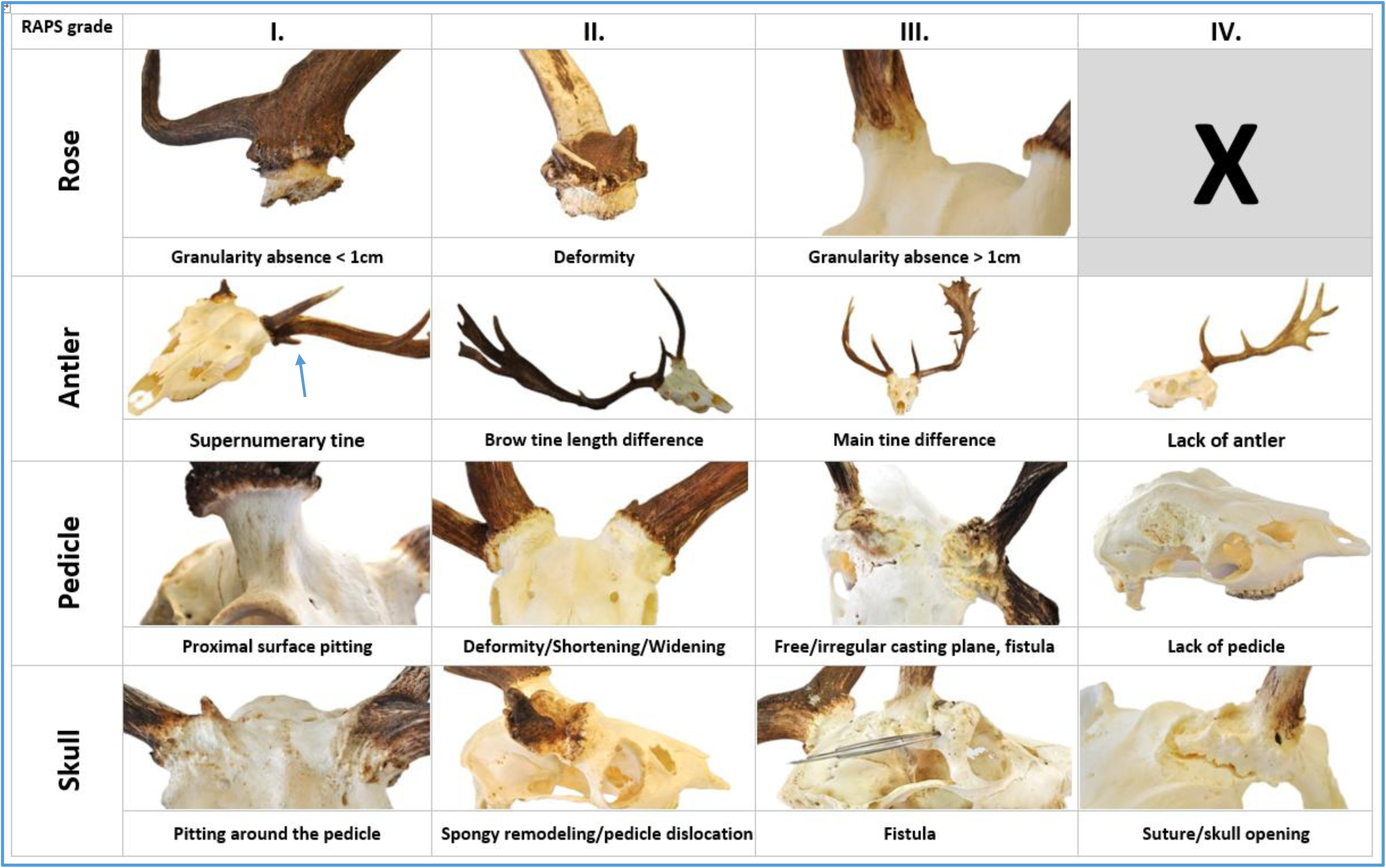
The recognition of the RAPS (Rose/Pedicle/Antler/Skull) features allows describing the severity of Pedunculitis Chronica Deformans (PCD). The RAPS features can be grouped into three grades for rose and four-four for pedicle/antler/skull. **Grade I.** Lack of continuity in the grain of the rose is longer than 5 mm but less than 1 cm. **(R1).** Supernumerary tine originating from rose, can be a sign of extra antler bud **(A1).** Pitting on the proximal surface of pedicle **(P1).** The pitting extends on the skull surface **(S1)**. **Grade II.** A deformity of the rose that is visible to the eye is a deviation from a regular circle. This is not independent of the pedicle deformity (P2) and visible on shedded antlers, too **(R2).** The length difference between the two sides of the brow tines **(A2)** and difference from Trophy Register Control group **(A2.1)** which is derived from data of Hungarian Trophy Register after exclusions of cases marked abnormal (see text). The pedicle deformity characteristic to PCD in Fallow deer (Dama dama) is the deviation from the regular circle diameter more than 4 mm, also visible with eye **(P2).** Shortening **(P3)** and widening **(P4)** of the pedicle beyond normal extent. Spongy remodelation of skull bones where the cavities are larger than at intertrabecular spaces in spongious bone **(S2)** or visible the center of the pedicle has shifted significantly, mostly towards the brow ridge **(S6). Grade III.** Loss of rose granularity is greater than 1 cm **(R3)**. Our study shows that the R2 and R3 can also informative for subcutaneous lesions. The difference in length of the main beams relative to each other **(A3)** and the difference relative to the Trophy Register Control group **(A3.1).** A peduncle fistula **(P5)** (not visible) or a free casting plane where does not protrud any antler-bony protuberance **(P6)**. A fistula mouth opens distant from the pedicle on the skull **(S4). Grade IV.** Absence of antlers or aberrant antler beams **(A4).** More than fifty percent of the casting plane of pedicle is in or below the plane of the skull, considered to be the absence of it **(P7).** Discohesion along the interosseous suture **(S3)** or separation of cortical bone independently a sign of bone displacement **(S4).**

We compared the frequency of RAPS features between a healthy Morphological Control Group (MCG) and aberrant-antlered DD cohorts. On the MCG (24 trophies) from the Northern control area, we inspected 1,056 morphological features, of which 16 (1.52%) showed anomalies. In contrast, among aberrant DD trophies from our study area (50 trophies, 2200 inspected features), 989 pathological features were detected (44.95%) **(Supplementary Data 5)**. The Fisher’s exact test (p < 0.0001) confirmed that the RAPS characteristics were highly discriminative between these groups **(Fig. 5A)**. All of the RAPS features showed a consistently higher prevalence in the aberrant trophies than in the MCG, with 20 of the 22 being statistically significant discriminators (Pearson’s Chi-square tests, see **Supplementary Data 5**).

**Fig. 5A.**
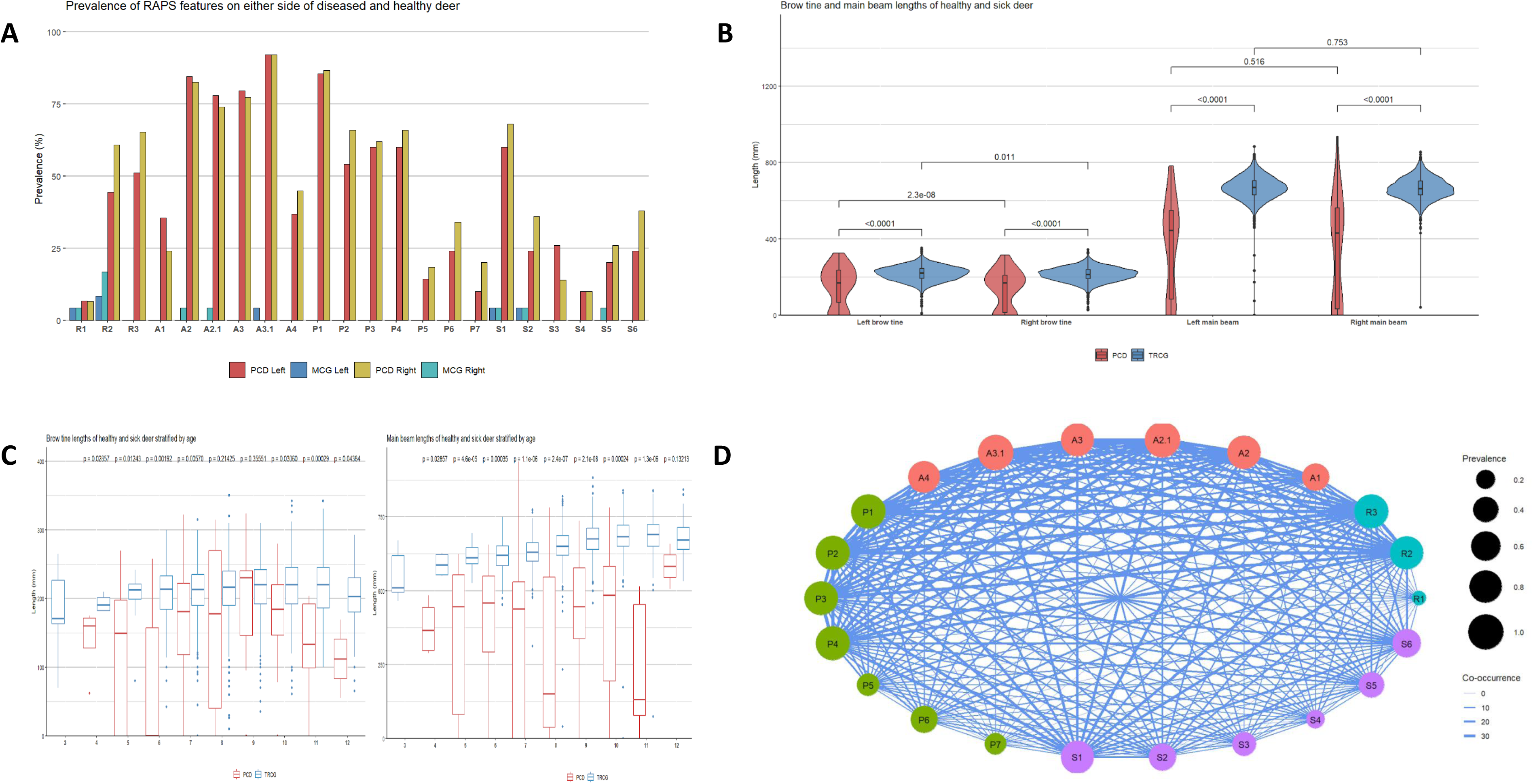
Frequency of RAPS features representing anomalies of the Rose (R) Antler (A) Pedicle (P) Skull (S) in fallow deer (*Dama dama*). 50 trophies with antler abnormalities (left side: red and right side: yellow) and 24 trophies from a healthy (morphological) control group (left side:blue and right side: green) were evaluated. Some anomalies were observed in healthy control individuals, but at low fequence **(Supplementray Data 5). B** The longness differences between anomalous DD (red) and healthy (blue) brow tinies and main beams by sides in comparison with Trophy Register Control Group (TRCG). There are no significant side differences in either the anomalous or the TRCG cohort. However, there is a significant difference between the two groups in both main beam and brow tine length. **C** Difference in brow tinies and main branch length between abnormal DD individuals and TRCG by age group. **D** The graph shows the co-occurrence of RAPS characteristics. The size of the nodes represents the prevalence of the symptom in the entire cohort and the width of the edges represents the number of co-occurrences between the two symptoms. The data are not aggregated by site. However, not all RAPS characteristics independent of each other may be able to describe more accurately the progression of this antler abnormalities. and to elicit a consensus based on a larger number of cases. The numerical data is visible in **Supplementary Data /**

While antler abnormalities are generally considered unilateral lesions, our study suggests a bilateral condition (**Supplementary Data 5)** arising from a systemic issue affecting the entire organism, as was previously suggested by Rachlow et al. and Kierdorf et al.^7,30^. The noticeable asymmetry likely reflects varying environmental factors, primarily infections and secondary traumas, as previously suggested^30^. This chronic process, which can span several years and multiple antler shedding cycles, becomes more pronounced on the side that is more severely affected.

In our study, the most frequently observed anomaly was— compared to the TRCG—a reduction in antler main beam length (A3.1, 92%) **(Fig. 5 B C)**, suggesting that the diminished bone-forming potential of the main beam bud cells is a sensitive marker for this condition. Additionally, visible asymmetry between the main beam length was frequently noted (A3, 69%), further supporting this idea.

The second most frequent change was the pedicle surface pitting (P.1, 80%), particularly visible on the frontal area, described earlier by Davidson et al. as a hallmark of this anomaly^31^. Histological and radiological analyses revealed that this accompanies the superficial form of chronic osteomyelitis ^32–34^ **(Fig. 6 d and e**). This often extended to the adjacent skull (S1, 64%). The rose deformity (R2, 48%) and discontinuity of rose granularity greater than 1 cm (R3, 53%) were also prevalent, appearing in roughly half of the cases. A prior study has noted the association of R2 with unilateral antler deficiencies^30^. In our study, R2 was significantly associated with visible brow tine difference (A2, 76%, while R3 correlated with shortened brow tine (A2.1, 76%) and main beam (A3.1) length. Furthermore, both R2 and R3 were significantly associated with all pedicle and skull anomalies **(Fig. 5C and Supplementary Data 6),** indicating that the more severe rose abnormalities can act as an indicator of subcutaneous issues. The irregular, also called “dirty” casting plane (P6, 29%) in shed antlers is a well-documented phenomenon^1^, associated with an insufficient testosterone peak^35^. However, only a third (29%) of the trophies displayed this characteristic, as many were collected outside the casting period. The shortening and widening of the pedicle is a physiological event that occurs with age^1,36^. Based on subjective judgment, shortening (P3) was considered more pronounced than normal in 61% of abnormal cases and widening (P4) in 63%. However, the pedicle deformity (P2), defined as a deviation from a regular circle shape appeared in over half of our cases (60%), serving as a, usable marker of disease recognition in DD but not in CC and CE **(Supplementary Data 1 and 2)**. In extreme cases, lateral displacement of the remaining pedicle, often extending to the supraorbital ridge (S6, 31%) was observed, similarly to the phenomenon photographically documented in North Arizona elk *(Cervus canadensis)* populations by J.L. Rachlow^7^. These aberrant proliferation-remodelling features (P2 and S6) are similar to those described in our study, suggesting that it is not just a DD-specific phenomenon.

**Fig. 6.**
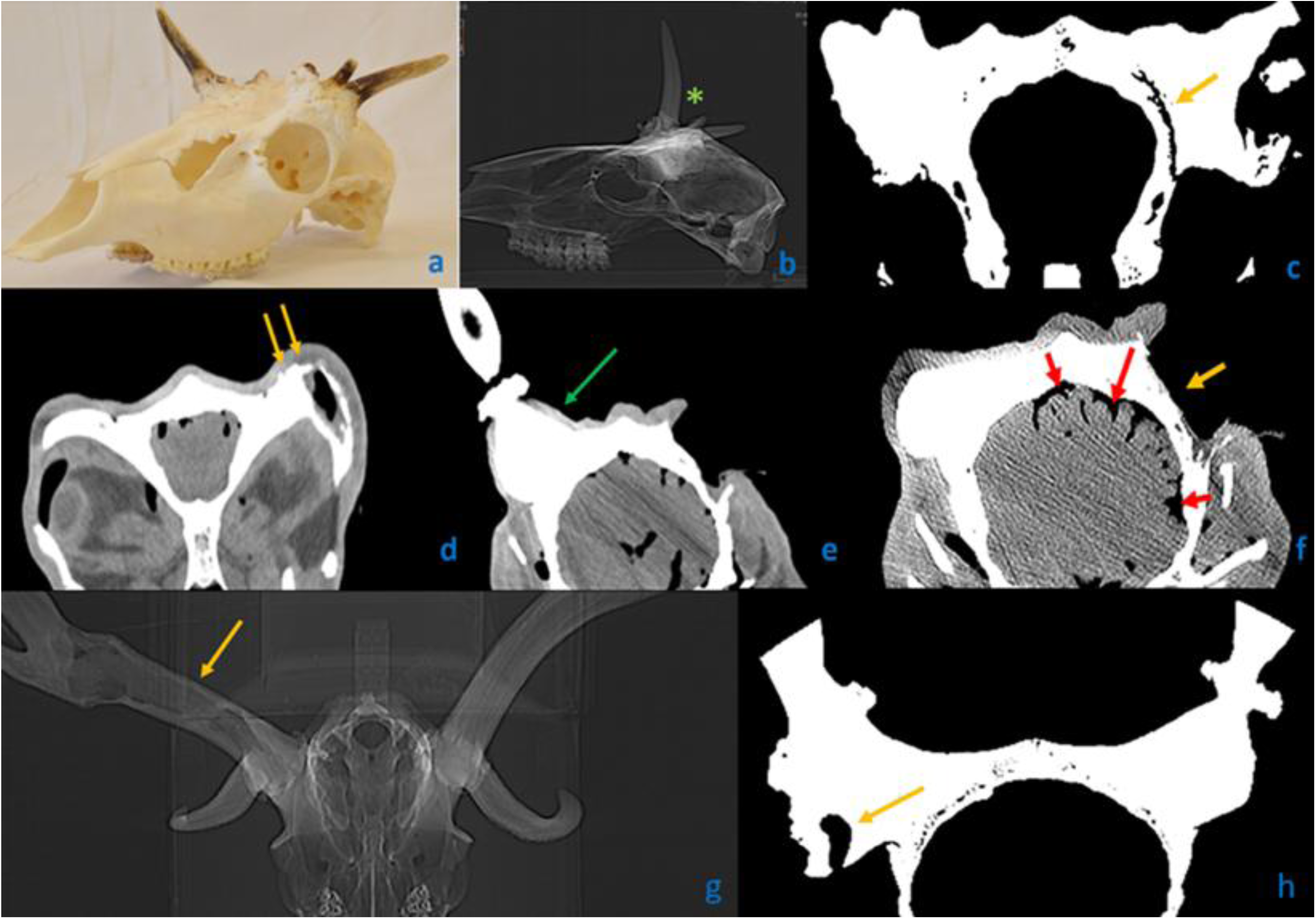
Computed tomography (CT) scans on three Fallow deers (Dama dama) cases. **a-c** A trophy with severe involvement on both sides with aberrant antler beams (AAB). **d-e** A complet head with soft tissues. **g-h** The only case with cavity formation in antler**. a** Bilateral loss of antlers with deformed peduncles and on them AAB**. b**Typically, no cavity is formed in aberrant antler beams (marked with green asterisk)**. c** Frontal scan through antler pedicles. On the left side, not visible to the naked eye, a large gap-like cavity (yellow arrow) running parallel to the cranial plane without displacement of the cranial bones. Its development due to trauma is unlikely. **d** The whole head with soft tissues. The yellow arrows show the unevenness of the residual peduncle suggestive of superficial chronic osteomyelitis. **e** The green arrow shows in the deeper plane the other side peduncle surface without significant radiographic abnormality. **f** In an even deeper plane due to pus accumulation in the subcranial space the widening of the sulci and thinning of the gyri (red arrows) suggest the meningoencephalitis. Yellow arrow shows the lack of peduncle**. g** Only one of our cases showed a cavity (yellow arrow) in the unilateral antler beam, which may indicate the death of central blastema cells during growth**. h** The necrotic cells may have been exited through a fistula (yellow arrow) to outside.

In our work, we defined “spikes” (a term used within the hunting community) as *Aberrant Antler Beams* (AAB) to distinguish terminologically between diseased forms and healthy one-year-old bucks without branched antlers. If nothing or only an AAB was present at a given site, we classified the specimen as “lacking antler” (A4, 40%). We considered a formation as an antler if the main beam exceeded the brow tine in length, with at least a rudimentary antler rose. CT scans revealed these structures to be generally solid, with only one case showing cavity formation **(Fig. 6 g-h)**. In cases where pedicle blastemas are less involved, more pronounced growth can occur by incorporating unused nutrients and minerals, which become more available due to a reduced number of osteoblasts in other areas. We termed this process *Compensatory Growth* (CG). The effect might be analogous to contralateral antler growth following removal of the peripeduncular periosteum described by Li et al^37^. CG can be indicated by a significant difference in brow tine length due to a longer-than normal brow tine on the healthy side. The discrepancy can arise from shortened brow tines on the affected side due to damage of the local blastema, while the CG on the contralateral side results in a longer brow tine.

Among the observed skull abnormalities, irregularities such as peripeduncular skull surface pitting^31^ (S1, 64%), spongious skull remodeling (S2, 30%), and the lateral displacement of the expanding pedicle (S6) suggest atypical resorption and proliferative processes cannot be solely attributed to trauma or displacement during healing. A morphological evidence of secondary healing of chronic osteomyelitis is fistula formation ^32,38^, which was detected on the pedicle (P5) in 16% of the cases and on the skull (S4) in 10% of the cases. The opening of sutures (S3, 20%) theoretically might be a consequence of injury; however, it would not occur in isolation since it typically requires adjacent skull fractures and dislocations. Trauma-induced extensive detachment of the external cortical plate from the underlying bone layer would result in substantial displacement. Conversely, CT imaging **(Fig. 6 c)** of an aberrant trophy revealed that the gaps between cortical plates (S5, 23%) often aligned with the cranial contour, suggesting that the origin of this malformation is unrelated to trauma. The absence of any signs of sharp fracture lines—either with or without displacement—and the lack of callus formation or (at least) partially broken pedicle structures (f.e., regular circular cross-section, with a missing part) further support that these abnormalities are not typical traumatic injuries. If not only trophies are available but also soft tissues can be examined, the presence of abscess-like purulent inflammation is the evidence of the underlying pathology **(Fig. 7d-f).**

**Fig. 7.**
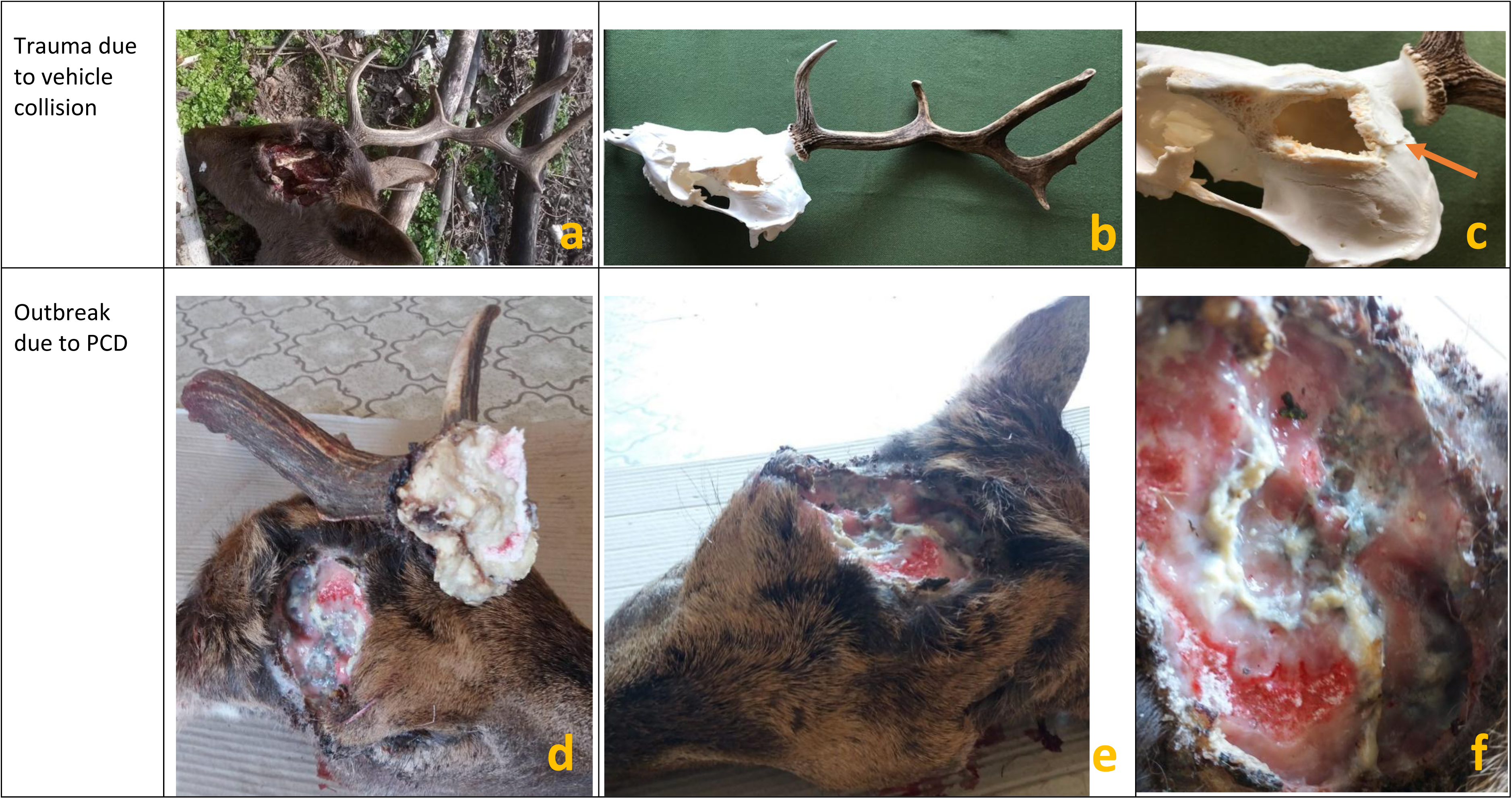
The difference between an obvious trauma caused **(a-c)** and a pathological fracture **(c-f)**. **a** The top row of images shows the left antler broken-out following a motor vehicle collision. **b** The condition after the soft tissue removal. **c** The fracture surfaces are sharp and angular, with displacement of the cranial bone (arrow). **d** Bottom row shows the right antler and the site of the broken-out, which happened due to a fall following a hunting gunshot. The surface of the break-out, which covers a large area, is rounded. **e** The loss of the left antler occurred earlier. **f** Most of the outbreak surface shows a greyish-white pus, indicative of preexisting osteomyelitis.

The prevailing hypothesis suggests that post-traumatic deformity within antler pedicles is a consequence of the misalignment of the broken surfaces since the periosteum and skin cannot hold them together properly ^1,30^. Certain studies seemingly supported this theory by examining the mechanical disruption of the pedicle immediately after casting or of antlers during the velvet stage^1^. However, it is important to note that these experiments were limited to growing antlers, which may represent imperfect antler growth rather than the process of pathological healing after the breakout of dry antlers ^39,40^. In our study, we examined one case where injury occurred during the velvet stage, which caused acute osteomyelitis, accompanied by re-epithelialization of the exposed velvet surface **(Fig. 8)**. This inflammation was localized within the injured area and its immediate vicinity, sparing the deeper marrow area in larger tines and avoiding extension to the pedicle. Consequently, healing produced a deformed antler with a size reduction proportional to the initial injury. In theory, a severe acute inflammation that reaches the pedicle could cause a fatal sepsis. A less severe process, confined to the antlers, can cause deformation or breakage of branches ^30,41^. However, animals mostly avoid injury during the velvet stage due to the innervated and sensitive nature of their antlers ^1,30^. In the absence of chronic inflammation or bone remodeling, injuries at the velvet stage are not usually a significant cause of antler deformities; healthy antlers can regrow in the following year ^1^. Our biobank reflects only one such case among other abnormal samples.

**Fig. 8.**
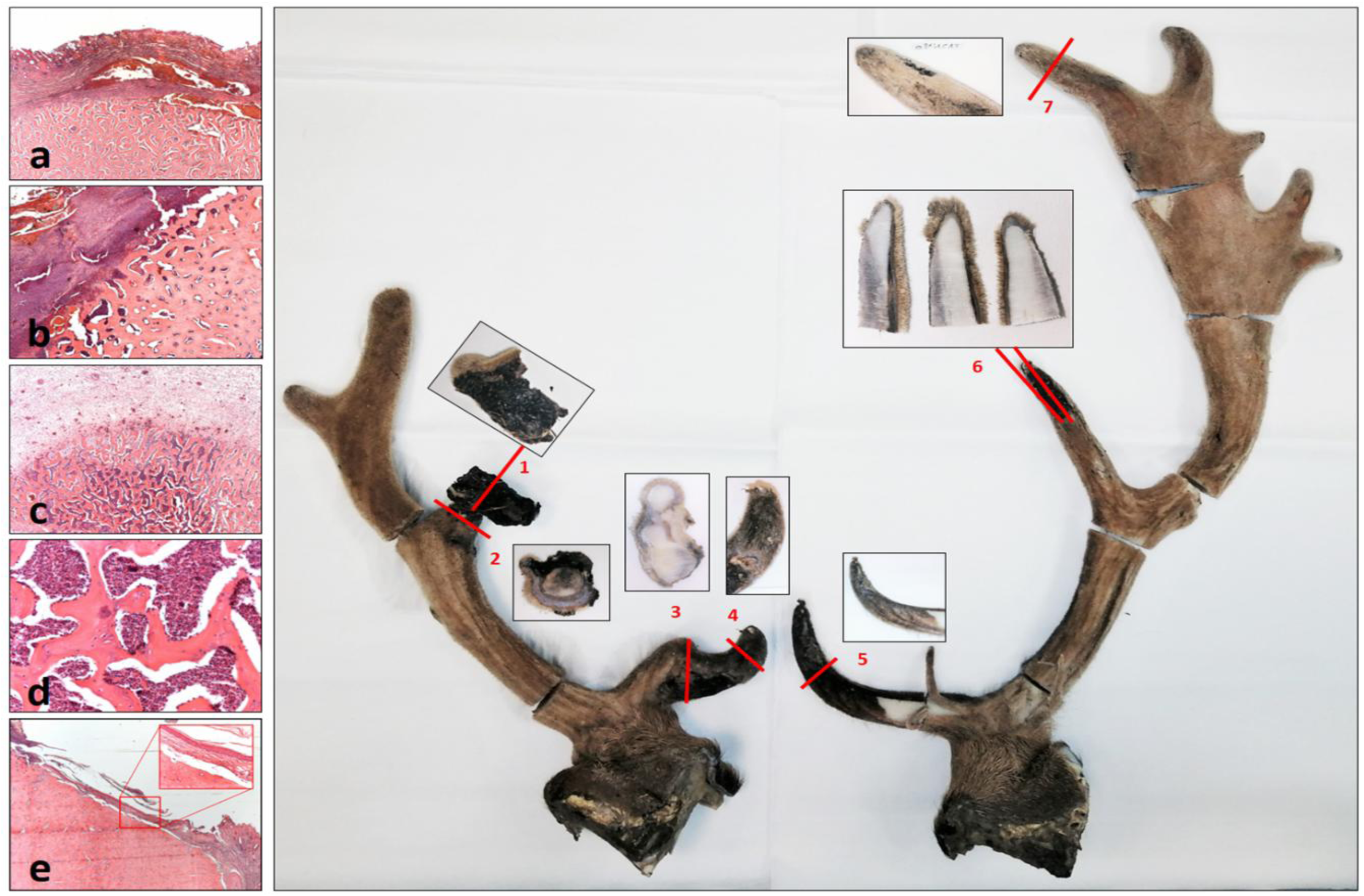
Microscopic examination of wounds caused by injury in velvet phase leading to antler deformity and asymmetry suggests acute inflammation. The significant bilateral difference may be related to the different extent of injuries and subsequent inflammation, and may indicative of a longer recovery time under while the antler could grow. The red numbers show the directions of the cut-offs for microscopic examination, the adjacent images show the cut surfaces of the planes. **a** Subcutaneous haemorrhage and detachment from the bone surface is an indication of injury (Cut 7). **b** The larger lesion is associated with necrosis of the skin and periosteum (upper left half of the figure). The superficial infiltrated blood layer confirms that this was caused by trauma (Cut 2). **c** Inflammatory cells reach the deep spongious bone tissue, too (Cut 5). **d** The intertrabecular spaces of the spongious bone are filled with acute inflammatory cells (neutrophil granulocytes) (Cut 5). **e** During reepithelialisation, the epithelium slides from the skin surface adjacent to the lesion to the free bone surface. The resulting contact is consistent with one of the histological components described in peduncolo dermal junction (PDJ) and is also indicative of a longer period of regeneration. The insert shows the bony-epithelial contact without the dermis. The gap is a retraction artefact formed during fixation (Cut 6).

### Multimycotoxicosis, hormonal abnormalities and their relation to bone formation

In the absence of established animal or cell culture models specifically for investigating the impact of mycotoxins on antler development and regeneration, we utilized comparative research on bone development, cellular regeneration, and immune function from other species as proxies. A recent review has summarized the knowledge on the molecular mechanism through which mycotoxins impact long bone growth^42^. To investigate the potential link between antler abnormalities and mycotoxins (MTs) exposure, we measured concentrations of key MTs, including aflatoxins (AFs), zearalenone (ZEA), deoxynivalenol (DON), and fumonisin B1 (FB1). These analyses were conducted on the blood serum, liver, and muscle tissues of affected DDs and healthy control animals **(Fig. 9 and Supplementary Data 9)**.

**Fig. 9.**
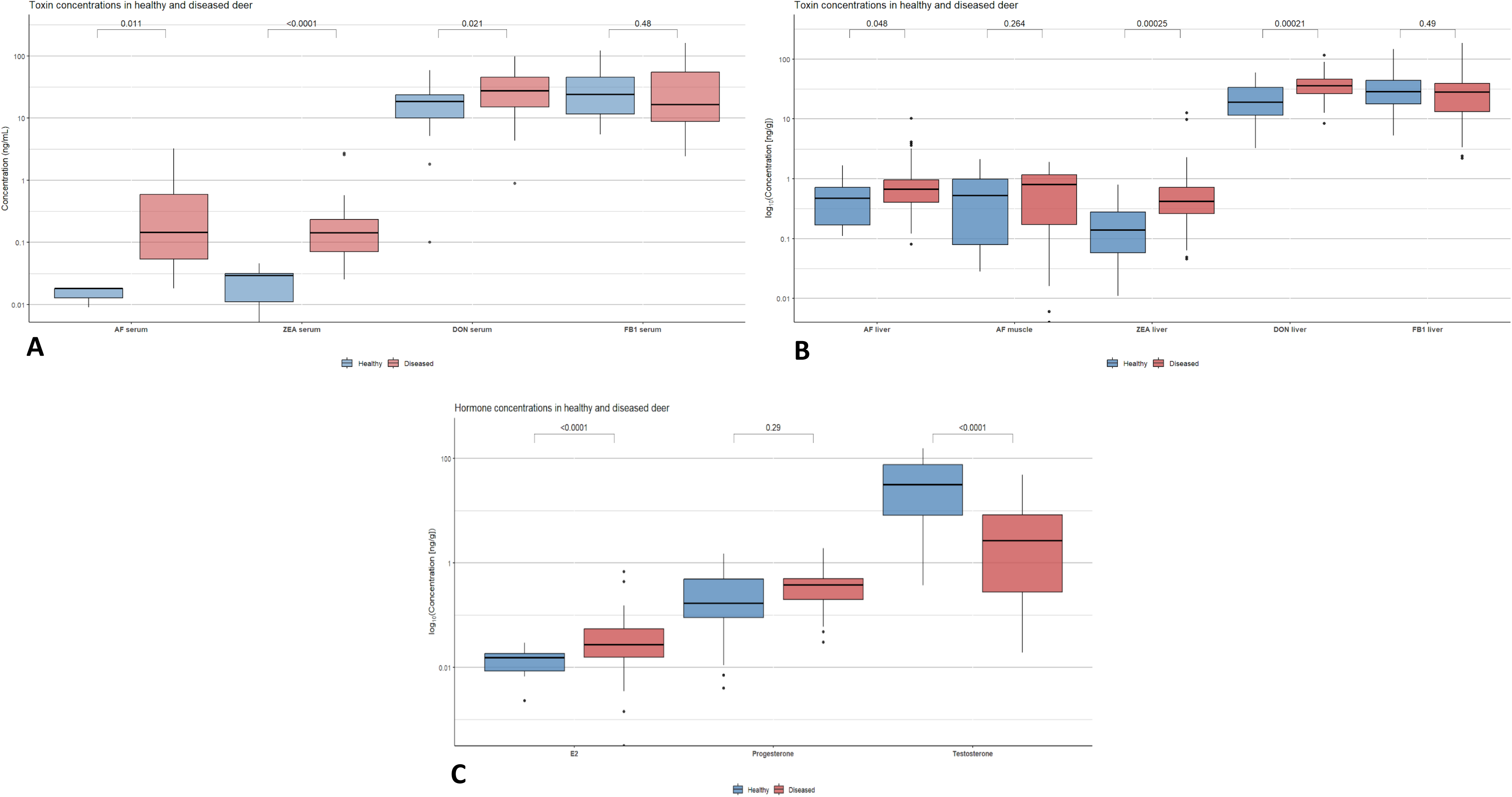
Mycotoxins (MTs) and hormons concentration of fallow deers (Dama dama) serum and liver in healthy and in the Pedunculitis Chronica Deformans (sick) cohorts. **A** All the MTs, Aflatoxins (AFs), Zearalenone (ZEA), Deoxynivalenol (DON) concentration were eleveted significantly in serum, except the Fumonisin B1 (FB1). The high level of AFs are responsible through direct cellular proliferation inhibition for the disruption of enchondral cartilage formation and addition with its immunomodulatory effect, for the inhibition of the cast wound healing. **B** All mycotoxins, except FB1, accumulated significantly in the liver. No increase in AF levels was observed in muscle. **C** Testosterone, progesterone and estrogen (E2) levels in serum. The significant decrease in testosterone levels can be attributed to the mycoestrogenic ZEA level increases. The elevation of endogenous estrogen levels can be the result of the compensatory activity of the hypothalamo-pituitary-gonadal system which, together with mycoestrogenic effect, may put double pressure on androgen-dependent antler development. The progesteron level not changed.

Our analysis revealed significantly elevated AF concentrations in serum (p= 0.011) and liver tissue (p= 0.048), with no significant increase in muscle tissue (p = 0.264) **(Fig. 9)**. This pattern aligns with AF’s mechanism of action in disrupting bone metabolism, specifically by inhibiting protein synthesis and impairing bone cell function^43^. A study employing micro-computed tomography demonstrated that AF exposure results in decreased bone mineral density and an increased risk of fractures. The effects of aflatoxin B1 (AFB1), a subtype of AF, are concentration-dependent and impact both bone health and nutrient absorption. At low exposure levels (75 ppb), AFB1 downregulates the jejunal mRNA expression of the vitamin D receptor, as well as calcium and phosphorus transporters, thereby impairing nutrient absorption critical for maintaining bone health. At moderate exposure levels (225 ppb), AFB1 reduces cortical bone volume in the femur metaphysis of broiler chickens. At high exposure levels (450 ppb), AFB1 significantly decreases trabecular bone mineral content and density^44^. This evidence suggests that AFB1 may disrupt antler formation by reducing the viability and differentiation potential of antlerogenic stem cells (ASC), which are highly productive and responsible for substantial tissue growth, generating approximately 3.3 million cells annually and contributing to the formation of up to 15 kg of antler mass each year^45^. Our research also highlights that AFB1 can impair this process by reducing the number of stem cells and hindering their ability to differentiate due to its direct cell-damaging effects. AS cells also possess immunosuppressive capabilities^45^. Additionally, AFB1’s Janus-faced effects on immune modulation, which include increasing pro-inflammatory interferon gamma (IFN-γ) levels in natural killer (NK) cells and reducing anti-inflammatory interleukin-4 (IL-4), may predispose casting wounds to superinfection due to impaired immune regulation ^15,43^. Moreover, AFB1 inhibited antigen presentation by dendritic cells in pigs^46^. Given AFB1’s immunosuppressive characteristic in antigen presentation by dendritic cells, these properties likely impede effective wound healing in antler tissues.

Our measurements identified significantly elevated ZEA levels in both serum (p<0.0001) and liver (p=0.00025) tissues **(Fig. 9)**, the latter being responsible for ZEA excretion. ZEA, a potent mycoestrogen with relatively low toxicity but has been shown to delay skeletal ossification, as evidenced by *in utero* studies conducted in rats^47^. Elevated ZEA intake has been linked to reproductive issues in cattle and sheep, and it is known to induce hyperestrogenic syndrome in pigs^16^. This mycotoxin exhibits antagonistic activity on androgen receptors and can disrupt androgen signaling pathways crucial for antler growth, which is androgen-dependent ^1,48^. One of our key findings was a significant decrease in testosterone (p<0.0001) and an increase in estrogen levels (p<0.0001) among cases of antler anomalies during the rutting season **(Fig. 9 and Supplementary Data 9)**. This phenomenon is likely attributable to the suppressive effect of ZEA on the hypothalamic-pituitary-gonadal axis. The irregular cast plane, which is the consequence of the insufficient testosterone acme^35^ serves as evidence of the connection between the effects of mycotoxins (MTs) and abnormalities in pedicle bone development. This hormonal disruption highlights serious reproductive and physiological consequences for deer populations exposed to ZEA. These findings underscore the need for further research into the endocrine system and *in vivo* antler stem cell models ^49^ to better understand ZEA’s impact on antler development and the broader systemic health implications for Cervidae exposed to MTs.

In affected animals, we measured serum DON levels approximately three times higher than normal (p=0.021), furthermore DON levels at liver tissue were also significantly elevated (p=0.00021)**(Fig. 9)**. DON is known to disrupt the delicate balance between bone formation and breakdown by inhibiting osteoblasts’ activity while stimulating osteoclasts^50^. This dual action leads to weakened bone structures and an increased risk of fractures. Moreover, DON exhibits immunomodulatory effects that are dose, exposure, time, and frequency-dependent^17^. These effects can significantly impact wound healing processes, especially in the context of the chronic conditions observed in affected *Cervidae*.

FB1 showed no significant increase in either serum (p=0.48) or liver tissue (p=0.49) **(Fig. 9)**.

We hypothesized that morphological evidence of the direct cytotoxic effects of mycotoxins on bone formation can be demonstrated by the cavitation observed in the pedicles of one-year-old bucks. At this developmental stage, antler shedding has not yet occurred, ensuring the external integument remains intact. Antler development follows four distinct ossification phases: intramembranous ossification, transitional ossification, pedicle ossification, and antler enchondral ossification^40^. MTs can traverse from the Fallow deer (*Dama dama*) hind circulation through the placenta into the fetal circulation and can also be detected in maternal milk^51^. In addition, according to the human studies they can be detected in the Follicular Fluid as well ^52^.

We propose that these mycotoxins not only inhibit the growth centers of antlers but also impede the enchondral ossifying tissue of the pedicle, leading to cavities filled with granulation tissue^53^. This results in lesions indicative of aseptic bone necrosis (ABN). ABN is recognized in humans as a primary, non-infectious bone necrosis, primarily affecting young, growing children. It manifests as focal necrosis beneath the articular cartilage surface, where resorption of the necrotic area creates a cavity, leading to surrounding bone collapse and the space becomes filled with granulation tissue ^54^. Factors like steroids, certain medications, alcohol and radiation are hypothesized to inhibit bone regeneration through prolonged chronic inflammation. Histologically, necrotic bone and marrow elements, along with fibrosis and neovascularization have been documented. Despite the limitations in microscopic examination of pedicle samples - such as desiccated samples with minimal residual tissue, lack of classical formalin fixation-evidence of necrosis and the presence of regenerative granulation tissue were observed **(Fig. 10)**. Unilateral presentation of these cases has been observed in three cases, which is attributed to the physiological antler asymmetry^1^ rather than environmental influences. One case in our *Cervus elaphus* (CE) aberrant trophy series can be attributed to the same pathomechanism: a three-year-old stag with a right pedicle fractured at its proximal level, while the distal portion and the antler remained largely unaffected. This observation suggests the presence of a pre-existing cavity. **(Supplement Fig. 2 A-D and Supplementary Table 2)**.

**Fig. 10.**
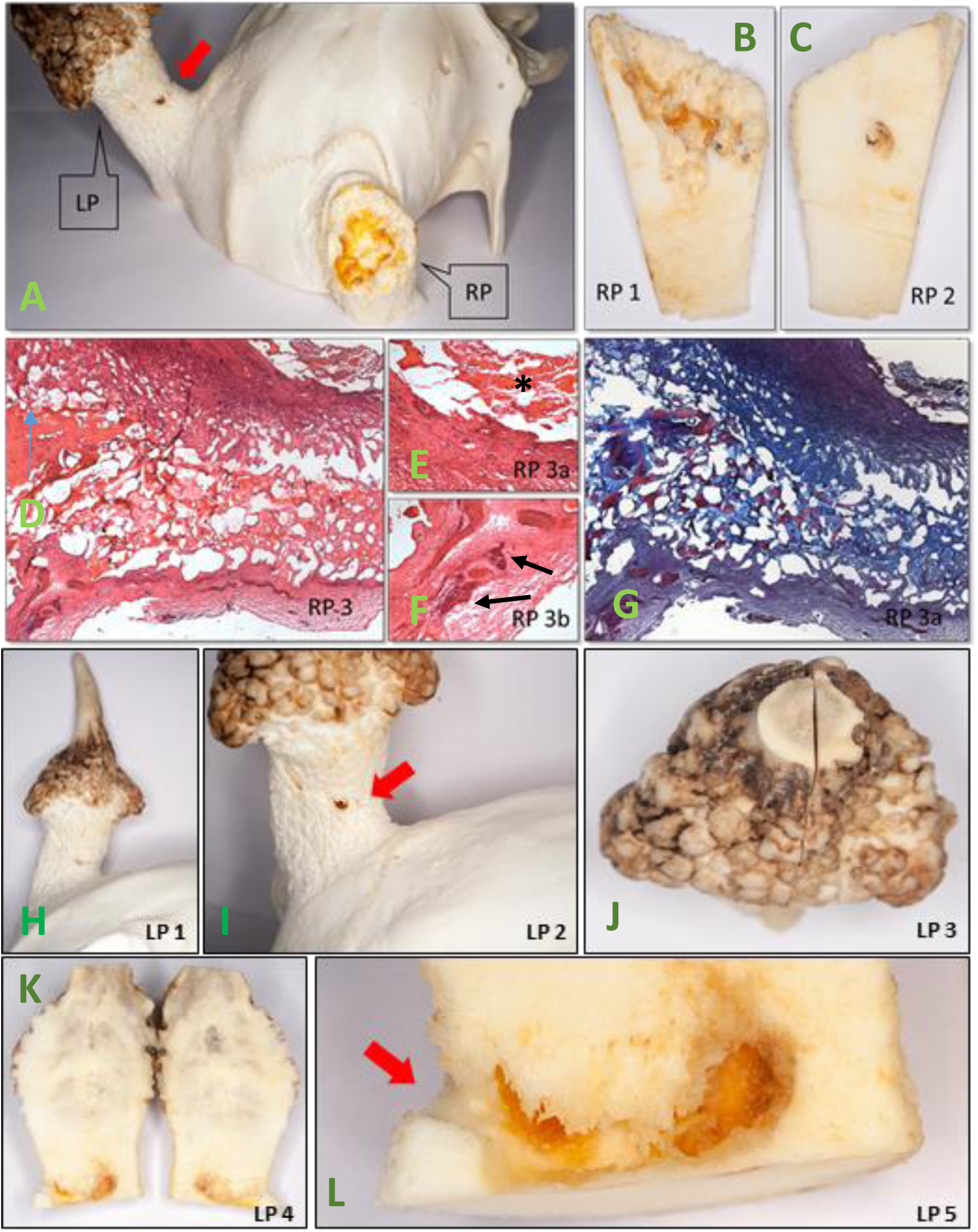
One-year-old stag trophy with aseptic bone necrosis (ABN) in the both pedicles. **A.** The right antler (RP) is broken-out and the fracture surface shows a partially lined cavity with a dry non-purulent exudate. The left peduncle (LP) has a pitted surface with a central fistula opening. **B** The longitudinal section of the right peduncle shows the extent of the cavity. **C** Parasagittal section with an early fistula formation. **D** Microscopic examination of the cavity base. Fibrotising granulation tissue with a subtle ossification is evident. (H&E 100x). **E** A higher magnification the D field upper right corner shows tissue detritus (asterisk) lined the inner surface of cavity (200x). **F** In a 200 times magnification the D field bottom left corner shows the perivascular accumulation of inflammatory cells (arrows). **G** Same microscopic field as G with Crossmon stain which is detect with bluish discoloration the fibrosis. The reddish spots indicative to osteoid material. **H** The left pedicle with the whole retarded in length one-year-old antler (spike). **I** The surface of the left peduncle (LP) shows pitting along its entire length, with a fistula (red arrow) opening in the middle**. J** The cutting plane made in the middle. **K** Large cavity formation deep below the casting plane in the area of early peduncular chondrogenesis is consistent with ABN. We suggest that, the region containing the rapidly proliferating cells was damaged by mycotoxin effect. **L** At higher magnification, visible the fistula opening to the outside, through which dead tissue could have exited, a well-known secondary phenomenon in abscesses, not a pathogenic portal of entry.

Given the direct negative effects of these MTs on chondro- and osteogenesis, they can also be considered *antlerotoxic*, adding new dimension to their well-known toxicities, which include hepato-, nephro-, neurotoxic, etc. effects. The pathological development of the pedicle along with the insufficient closure of the integument creates an environment that fosters bacterial superinfection, intracranial inflammation and pathological fractures of the pedicle and skull. This chain of events can ultimately lead to severe health consequences including the death of the animal.

### Healing of the casting wound

Upon casting, a wound corresponding to an open bone fracture forms at the detachment site. In healthy animals, this wound generally only leaves at most a small central scar after healing, getting eventually enclosed in velvet^25^. However, there is an ongoing debate about the specific mechanism of wound closure. One perspective suggests that full-thickness skin grows from the periphery toward the center, culminating in scar formation^37^. Alternatively, researchers propose that closure involves the formation of subepithelial granulation tissue with minimal surface fibrosis^25^.

This debate is particularly relevant, echoing Richard J. Goss’s assertion that “scars and blastemas are mutually exclusive”^1^. Acute wound healing proceeds through hemostasis, inflammation, cell proliferation, and tissue remodelling. While blood clotting is sometimes considered part of the inflammation stage ^55,56^, it is critical for initiating repair. In case of a casting wound, a blood clot forms over the wound where it is penetrated by vascular buds from surrounding dermal layer’s disrupted blood vessels at the fracture site, accompanied by macrophages and fibroblasts. Subsequently, this phase leads to granulation tissue formation, consisting of cell-dense, loose connective tissue, over which a new epithelial layer serves as the first line of immune defense. Over time, vascularity and cell density decrease while the collagen content of the extracellular matrix increases depending on the inflammatory response^56^. Complete wound closure with full-thickness skin is a carefully orchestrated, immune-controlled process involving fibroblasts migrating along endothelial buds^56^. When fully healed, the wound is sealed with both epidermis and dermis, representing its final state. Small areas of scarring from granulation tissue are often seen between the main beam and brow tine, typically where the velvet closes last and near the growth centers rather than above them ^25,56^. These minor scars generally do not interfere significantly with antler growth^37^.

Wound healing is a complex and delicate process that can sometimes falter, leading to chronic, non-healing wounds. Common initiating factors in such cases include infection, persistent inflammation, immunosuppression, and impaired circulation, often working together to hinder recovery^55^. Despite the regenerative potential observed in antler wounds, research has not yet focused on the presence of inflammatory cells—essential indicators of immune activity—or on the morphological traits associated with pathological healing in antlers. In a non-sterile environment, wounds are susceptible to microbial colonization. This poses a risk to the ossification zone as bacteria can spread through the bloodstream, potentially causing osteomyelitis^30^. However, when the immune system is robust and wound closure is timely, inflammation is generally minimal and self-limiting, reflecting a natural resistance to chronic inflammation and infection refer to the normal physiological conditions ^25^.

### The Pedunculitis Chronica Deformans (PCD) is a consequence of impaired wound healing

When tissue regeneration is impaired and immunosuppression occurs, in our cases due to multimycotoxicosis, wound healing disorders can develop. These conditions can trigger an inflammatory response that exceeds physiological levels, resulting in secondary or delayed wound healing, which is typically associated with excessive scar formation ^55^. Instead of forming a uniform, continuous layer, the resulting scar tissue is a braided, irregular structure composed of fibrous bundles ^57,58^. Such irregular scarring disrupts critical signaling pathways between the wound epidermis and the regenerating blastema^25^. The early chondrocytes, which originated from periosteal antlerogenic stem cells, form blastemas at the outer edges of the wound that develop into chondro-osseous columns during ossification ^26,59^. The morphological examination of abnormal pedicles and AABs indicated that these structures originate from sparse, scattered residual osteoblasts and osteo-chondroid trabeculae in the irregular scar tissue. The most obvious manifestation of this is the phenomenon described by Rachow et al. is the “spike-on-one-side”, (SOOS)^7^, but it is not exclusively unilateral(**Fig 11 A,B and Fig 6 a,b**). In this condition, the pedicle periosteum undergoes appositional intramembranous ossification but widens unevenly due to scarring, often extending laterally and even approaching the orbital ridge **(Fig 11 C)**. This abnormal growth pattern results in further shortening and more pronounced bone loss in the pedicle, which can ultimately lead to a “hummel” stage **(Fig. 11D and Fig. 12)**. The “semi-hummel” condition can also occur: In this case, there is an antler on one side showing minor deviations only, while on the other side it cannot develop **(Fig. 1 and Fig. 11E)**. In place of the missing pedicle, usually only a widened bone ridge remained, which did not correspond to the morphology of a secondarily healed fracture. These trophies are thought to correspond to cases in which, despite a significant bilateral weight difference, the animals are able to keep their heads upright (**Fig. 1 and Supplementary Fig. 1**). In these cases, there is enough time for the development of a compensatory head-hold with neck muscle hypertrophy during the period of gradual weight gain on the intact side, providing an adequate visual field. Sudden weight loss, observed during non-simultaneous shedding, is associated with an oblique head position ^23^. Unilateral antler loss due to trauma would cause the same.

**Fig. 11.**
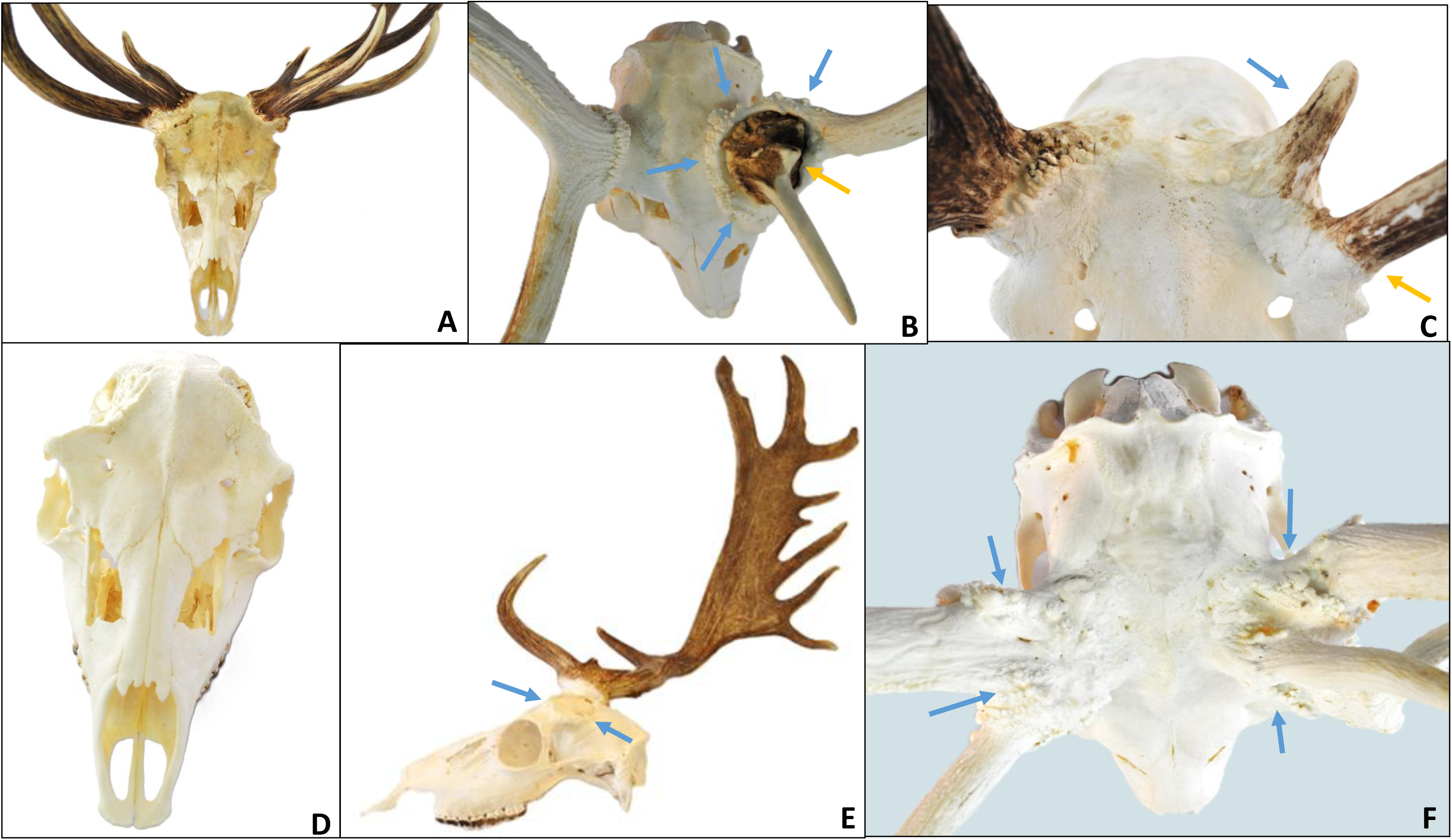
Morphological changes in Pedunculitis Chronica Deformans (PCD) suggest disrupted osteogenesis of the skull and antlers by scar tissue. **A** The bilateral aberrant antler beams (AAB) can be considered derivatives of the antler-forming buds that have retained their developmental capacity between the dissecting scar tissue. They correspond to the phenomenon described as spike-on-one-side (SOOS), but can also be bilateral. **B** Top view of a velvet phase trophy showing intramembranous ossification (blue arrows) around the left not casted AAB (yellow arrow). In this case, AAB causes lateralisation of ossification of antlerogenic stem cell origin chondro-osseous tissue. This process increases the diameter of the pedicle and also moves it laterally. **C** In cases of severe ASC loss, little residual antler tissue can form (blue arrow), typically on the periphery, often located on the orbital arch (yellow arrow). **D** Bilateral total peduncle loss leads to the „hummel” condition. **E** The place of left peduncle remained only a widened bone ridge like edge (blue arrows), which not correspond to a condition after a secondary healed fracture. The widening of the right peduncle is suggestive of PCD and the long brow tine and massive palm formation can be consider of compensatory growth. **F** The bony trophy could give a evidence to the scar formation indirectly only. However, its appearance with small nodules (blue arrows) corresponds to scars dissecting chondro-osseous tissue.

**Fig. 12.**
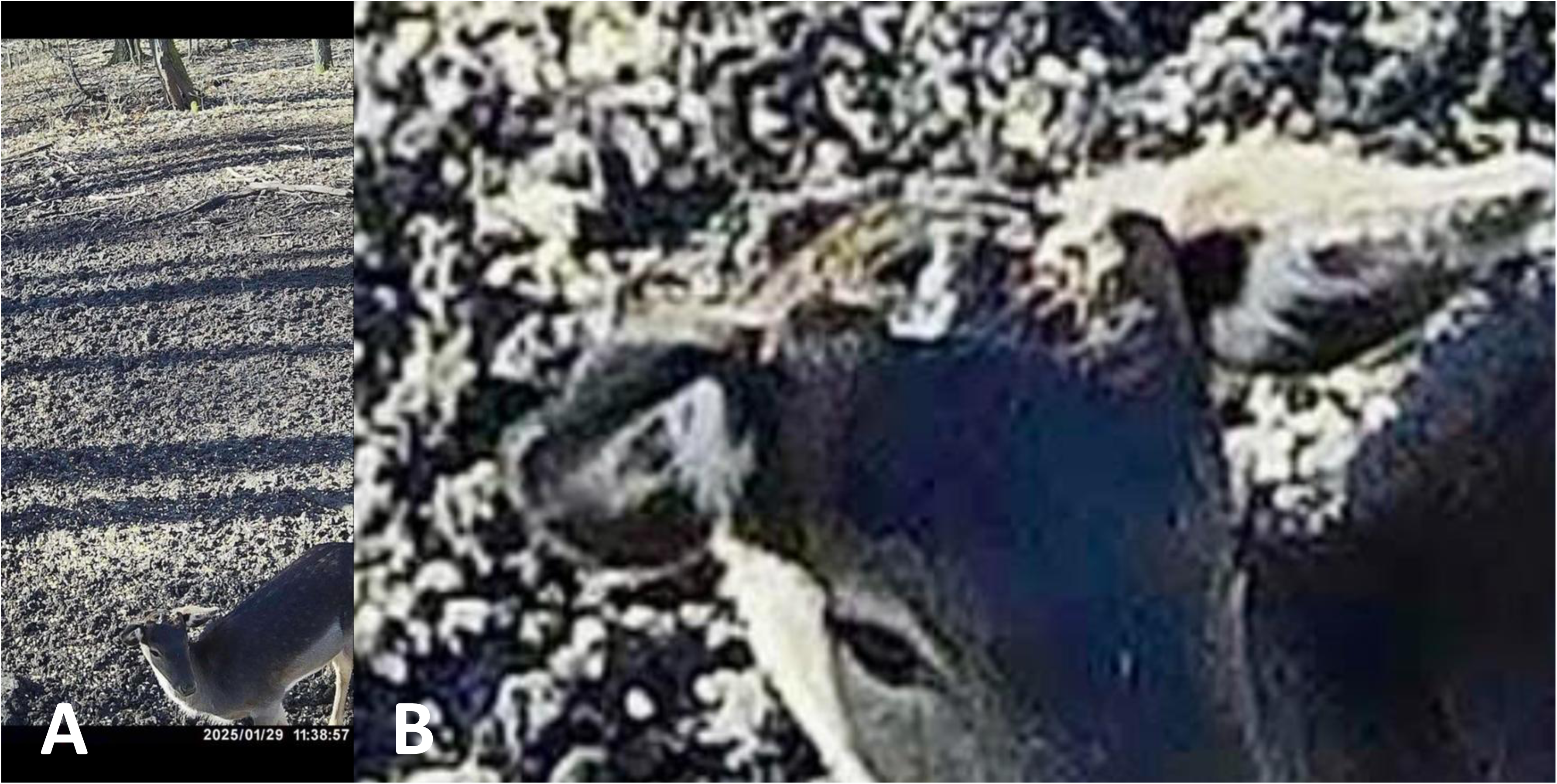
ATrail cam footage from around the village of Szakcs (South-Hungary, Middle Europe) from our study area. This living “hummel” fallow deer (*Dama dama*) buck proves that the losses of antlers on both sides is survivable. **B** However, the survival time is questionable. The magnification shows a rather large loss of skull fragments on the left side.

The etiology, pathomechanism, and morphology of the lesions we described do not align into any established classifications but share similarities with Garré’s sclerosing chronic non-suppurative osteomyelitis. This condition, which typically affects the mandible, is characterized by its “onion-skinning” bone deformation due to mild irritation or bacterial infection resulting from a proliferative periosteal reaction^60^. We have termed this unique condition *Pedunculitis Chronica Deformans* (PCD). In PCD, the osteoblasts of the peduncular surface ossify intramembranously but fail to form organized, parallel lamellar bone. Instead, they create irregular ridges derived from osseochondroid columns. The top of these ridges retains variable amounts of antlerogenic stem cells. PCD compromises the structural integrity of both the pedicle and the forming antler, occurring independently of any prior injury. The bony trophy can only provide indirect evidence of scarring. However, the appearance of small osseous nodules is consistent with scars that dissect bone tissue **(Fig. 11 F)**.

Antler casting is primarily dependent on osteoclast activity across the pedicle, with the greatest extent of bone resorption occurring at the casting plane. Under physiological conditions, this process does not result in complete regeneration; instead, the pedicle shortens while its diameter increases through appositional growth ^1,25^. In case of healthy casting, the surface of the pedicle remains slightly concave and prickly due to fractured trabeculae, but the pedicle itself retains a circular cross-section and relatively smooth exterior^1^. However, during PCD, there is a notable reduction both in osteoblasts, which are essential for rebuilding the tissues, and also in osteoclasts involved in antler shedding. The distribution of osteoclasts becomes irregular and no longer concentrates exclusively on the casting plane, leading to an uneven casting surface. This disrupted remodeling weakens the pedicle structurally, which can render it incapable of supporting the developing antler, leading to fractures under the antler’s weight even under velvet phase **(Fig 13)**. PCD is thus a primary contributor to pathological pedicle fractures. Differentiating whether the irregular shedding surface is a result of an atypical, remodeled casting plane or a pathological fracture requires close examination of the fracture surface and careful consideration of the timing relative to the antler shedding period. Previous work by U. Kierdorf and colleagues described an instance where a pathological condition was predisposed to the antler breakage following a velvet-stage injury, which compromised the antler but not the pedicle. They underscored that the underlying pathologic process is necessary in weakening the antler structure, thus predisposing it to fractures ^30^.

**Fig. 13.**
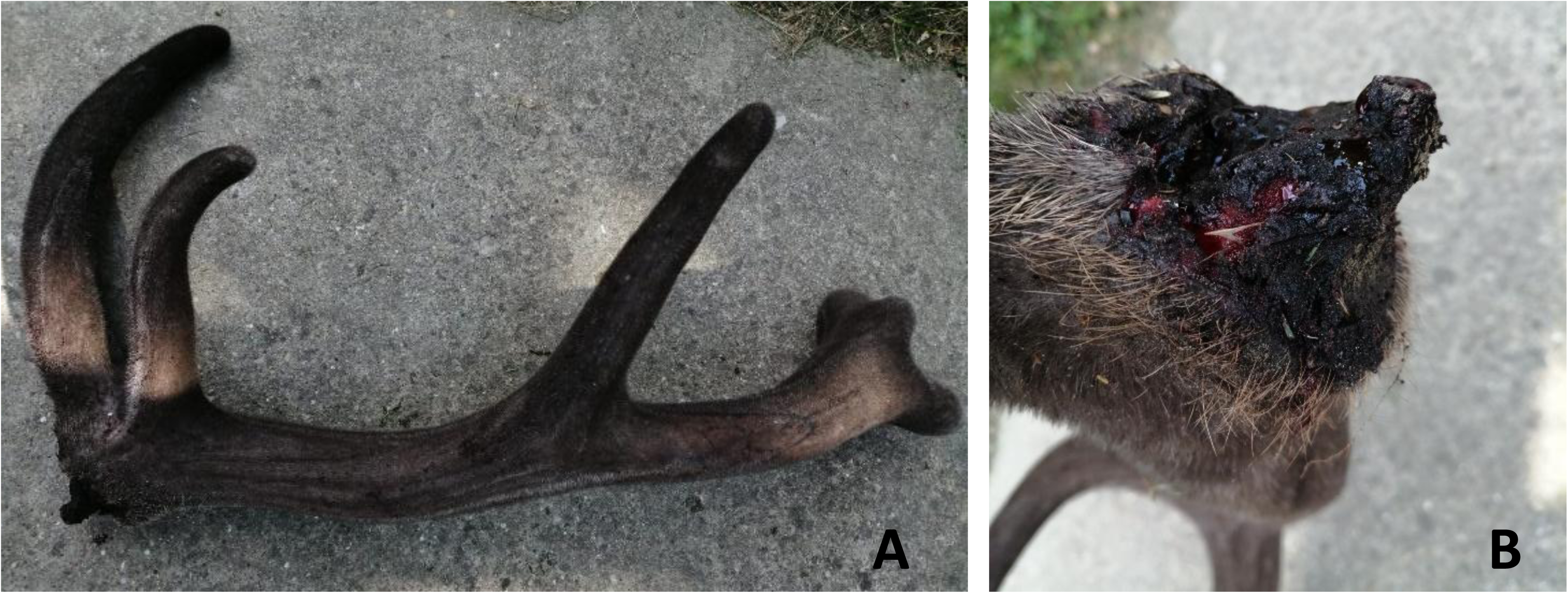
Antlers broken-out with an irregular plane in velvet phase. **A** There is no evidence of major force on the antler and it can therefore be assumed that the pathological fracture was due to the antler’s own weight. **B** The irregular fracture surface, not compatible with traumatic impact, was formed above the casting plane.

Thus, PCD can be defined as a chronic inflammation primarily affecting the pedicle tissue, caused by casting wound closure or enchondral ossification disturbance of the pedicle. The consequent scarring causes disturbance of pedicle and antler regeneration and weakening of their structure, creating the potential for pathological fracture, superinfection and the development of fatal intracranial inflammation. The PCD signs described by RAPS features are visible on trophy (**Supplementary Fig. 5**) and on harvested animals (**Supplementary Fig.6 and 8)**.

### Characteristics of pedicle infections and bacterial agents

Microscopical examination of the pedicle and surrounding soft tissues (skin, connective tissue, and muscle) revealed an acute inflammation. In deeper layers, chronic inflammatory cells and associated scar tissues were observed. This appeared as perivascular and perineural lymphoplasmacytic infiltration, with sequestra of detached bone from the pedicle surface embedded within the scar tissue—both characteristics consistent with chronic osteomyelitis **(Fig. 2i and 2m)**. No granulomatous inflammation or acid-fast bacilli indicative of infections like *Mycobacterium leprae* (the most common pathogen underlying this inflammatory pattern) were detected, as confirmed by negative Wade-Fite staining results. These histopathological findings across five examined cases were consistent with chronic periostitis, further complicated by acute secondary infection. PCD causes a condition known in hunting communities as “antler rot” through super-infection with pus-causing bacteria (**Supplementary. Fig. 8**).

This stage is accompanied by irregular bone remodelling closely resembling secondarily healed fractures. The infection has the potential to extend through the cranial bone leading to severe intracranial infections (ICI) such as meningoencephalitis and brain abscesses^31^. Our CT imaging of entire deer heads, including soft tissues, confirmed that inflammation surrounding the pedicle can indeed propagate to the intracranial space **(Fig. 6 f)**. Studies on North American white-tailed deer have reported the prevalence of intracranial infections ranging from 1.8% to 4% of the population^11,31^. These rates increase to approximately 9 % in bucks over three years old, highlighting intracranial infections as a non-negligible natural mortality factor in mature males^10^. These works found such a strong correlation between cranial bone deformities and ICI that it was suggested that bone lesions could be diagnostic indicators of possible brain involvement^31^. PCD encompasses a spectrum of bone abnormalities, where the most severe forms are prone to complications with ICI. Noteworthily, Davidson et al. identified cranial bone lesions, including erosion, suture separations, and pitting, that correspond to certain PCD symptoms, such as S1, S2, and S3. However, the likelihood of which bony lesions are associated with ICI requires further investigation to be truly informative.

*Trueperella pyogenes* (formerly known as *Arcanobacterium pyogenes*, *Actinomyces pyogenes*, and *Corynebacterium pyogenes*) is widely regarded as a primary pathogen implicated as accessing peripeduncular soft tissue inflammation ^10,11,31^. However a Hungarian case study also noted *Staphylococcus xylosus* as a potential pathogen ^61^. The presence of *T. pyogenes* among other microbes in intracranial suppurative meningoencephalitis disease further suggests its central role^62^. Consistent with these findings, our conventional culture analysis of 42 cases identified *T. pyogenes* in 24 (57%) *of cases* (**Fig. 14 A)** frequently coexisting with other microbes. Our next-generation sequencing (NGS) analysis, which is informative about the number of bacteria in the lesions too, further classified the microbial landscape of pedicle inflammation cases. *T. pyogenes* was detected in all six samples examined. However its relative abundance was notably low, reaching only 5.2% in one case and ranging from 0.1% to 1.5% in others **(Fig. 14 B and Supplementary Data 10)**. This, however, suggests that it is present as a contaminant rather than a dominant pathogen which is consistent with its ubiquitous, opportunistic nature. Typically residing on skin and mucosal surfaces in animals with an invasive potential. Following the minor skin injuries can colonize the host and necrotic tissue ^63,64^. Interestingly, *Fusobacterium* species emerged as the predominant bacteria in five of the six cases, comprising between 39% and 78% of the microbial population, while in the sixth case, it was the third most abundant genus at 19% **(Fig. 14 B and Supplementary Data 10)**. *Fusobacterium* species, obligatory anaerobic commensals, are often implicated in abscesses and osteomyelitis (as confirmed in human cases), especially when skin barriers are breached ^65,66^. The shift from traditional culture to NGS methods provides a fuller picture of pathogen prevalence and relative abundance, suggesting that *Fusobacteria* might be more relevant to ICI risk than previously understood through culture studies alone. It can be assumed that the media of classical culturing may provide a selective advantage for *T. pyogenes*, which is not the case of NGS. A deeper species-level analysis, particularly examining the role of *Fusobacterium nucleatum*, was beyond the scope of this study but could yield further insights into the mechanism of superinfection of PCD. However, both *T. pyogenes* and *Fusobacterium sp.* require damage to the integument to exert their abscess-forming effect ^31,65,67^. This is predicted by a significant difference in occurrence between the sexes in mating struggles ^31,62^. However, it is unlikely that these injuries only affect the immediate area around the pedicles. Among our study samples, we did not find any cases of abscesses or bony lesions adjacent to healthy pedicles on the skull that would suggest an inoculation effect caused by head injury. In our view, the inhibited healing of the casting wound opens the way for secondary infection-causing pyogenic bacteria. This is evidenced by the abscess in the pedicle, accompanied by purulent exudation and fistula formation around the antler as early as in the velvet phase (**Fig. 15)**.

**Fig. 14.**
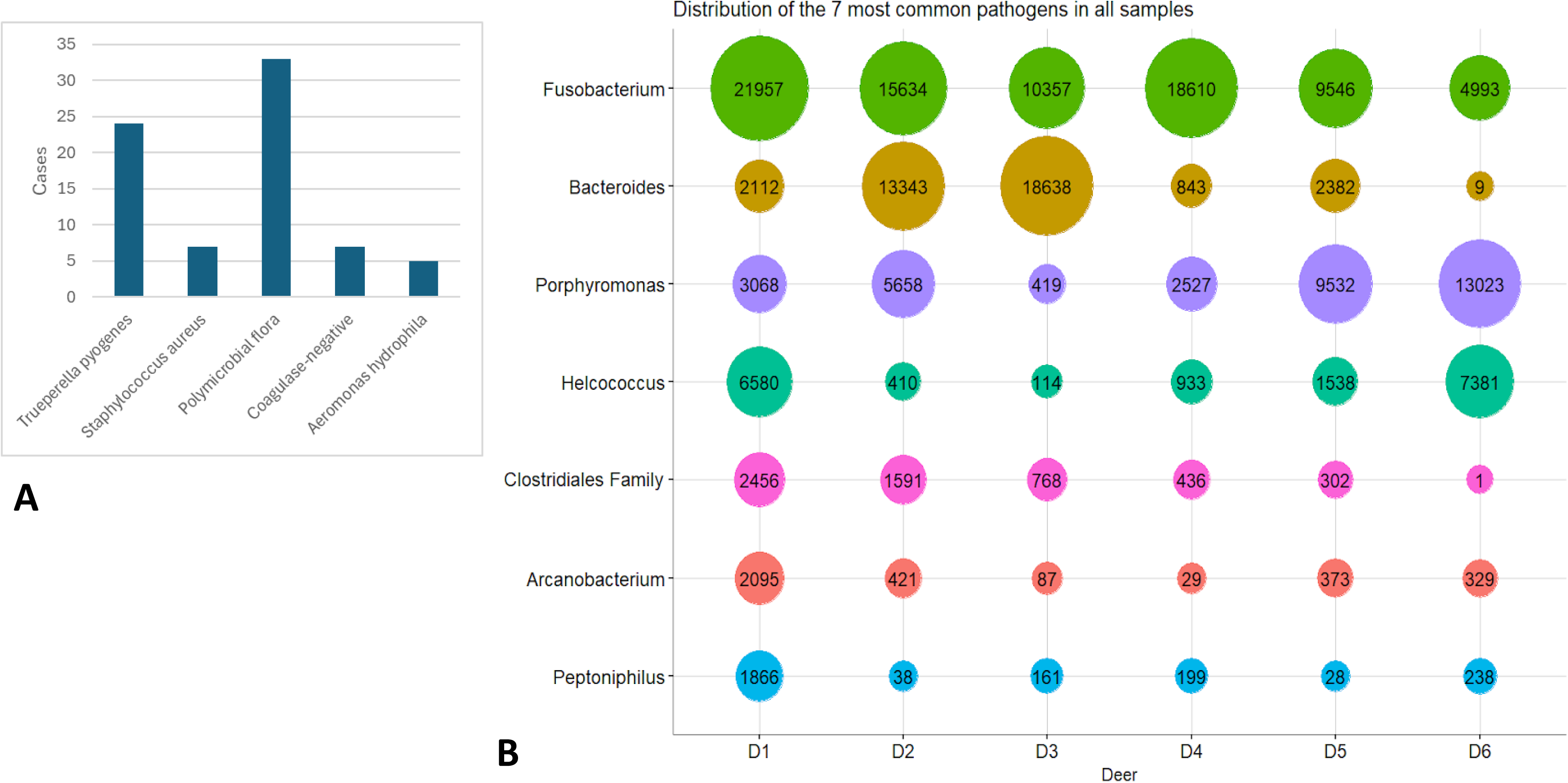
The microbiological examination of the peri peduncular inflammation. **A.** The conventional culture analysis of 42 cases identified *Trueperella pyogenes* in 24 (57%) of cases frequently coexisting with other microbes, which is consistent the results published by other authors (see text). **B** The next-generation sequencing (NGS) analysis, which is informative about the number of bacteria in the lesions too, further classified the microbial landscape of pedicle inflammation cases. *T. pyogenes* was detected in all six samples examined. However, its relative abundance was notably low, reaching only 5.2% in one case and ranging from 0.1% to 1.5% in others. In contrast, *Fusobacterium* was dominant in five out of seven cases but was the second and third largest bacterial colonies in the remaining two cases (D3 and D6).

**Fig. 15.**
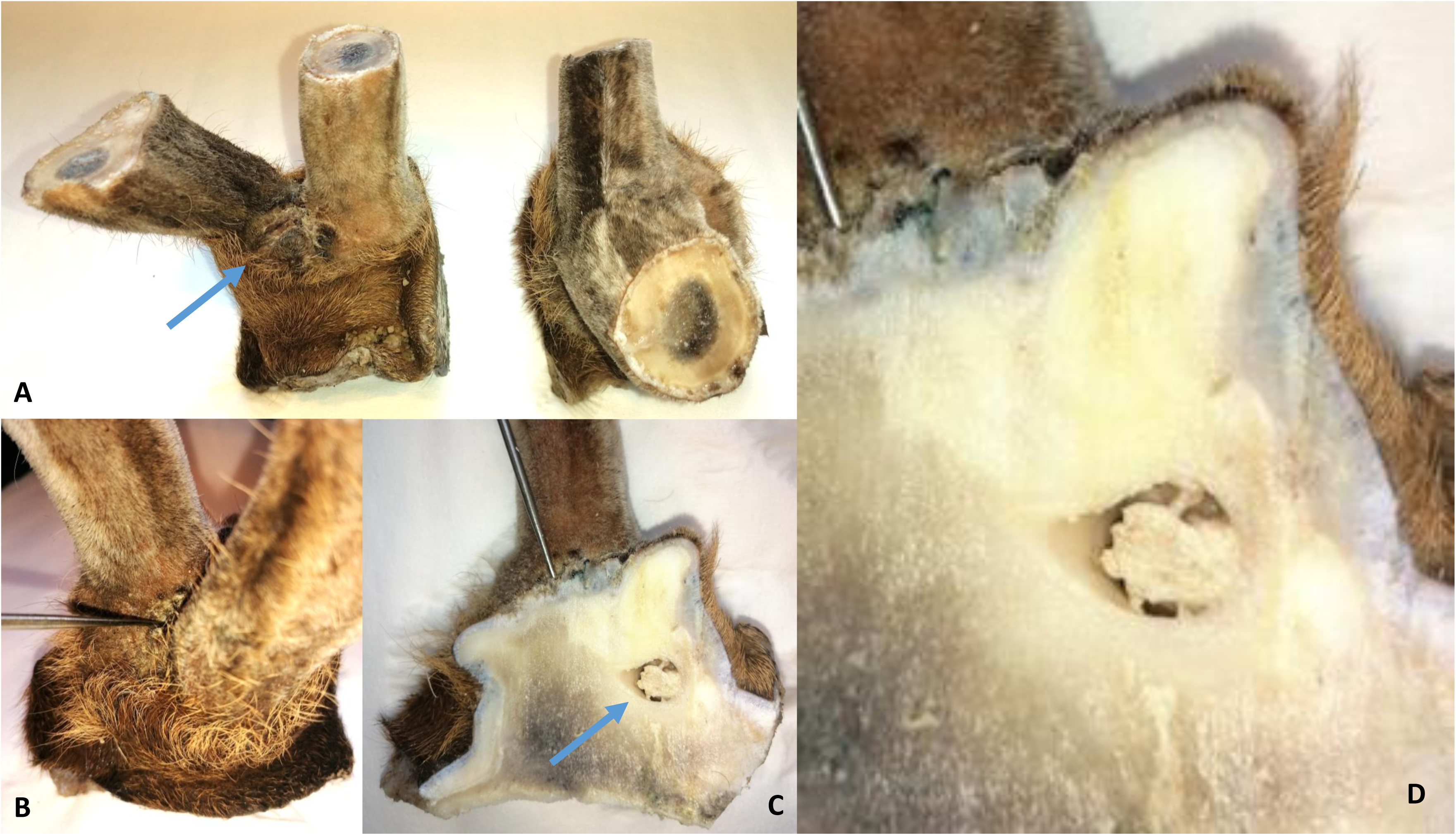
Abscess formation in the velvet antler pedicle. **A** On the left side, purulent peduncular exudate on the skin between antler beams indicates inflammation (arrow). **B** A fistula opening on the surface (metal probe in the fistula orifice). **C** An abscess cavity in the pedicle (arrow). **D** Note the peduncolo-dermal interface irregularity on the abscess closest surface of peduncle.

### Impact of global warming on the geographical distribution of MT concentration in Hungarian feed

Various factors may contribute to the development of antler and pedicle anomalies, including reduced mineral content in animal feed, changes in its chemical components, and behavioral shifts due to altered sex ratios ^2,7,68^. Initially, we suspected an infectious cause of the concentrations of PCD in southern Hungary, though no specific pathogen has been identified as the primary culprit. Our research highlights consistently elevated levels of MTs in livestock feed produced in Southern Hungary compared to Northern regions therefore we assumed it may play a role in the condition’s regional prevalence. AFB1 appearance in Hungarian maize kernels was first documented over 15 years ago^62^, with subsequent studies confirming the presence of DON and FB1, especially in Southern areas, including our study region, ^63^. This aligns with similar findings from Serbia^71^, just South of Hungary, underscoring a regional MT exposure pattern that could impact antler health.

Studies in nearby countries to the South of Hungary, including Croatia and Serbia, have reported a correlation between rising MT concentrations and changes in the local climate^72^. Weather conditions and AF production in our study area, as documented in previous research, were consistent with established patterns ^21,73^. The impact of climatic changes on MT production is complex, involving variables like temperature, radial heat, humidity, and the simultaneous effect of the wind, which collectively impact organisms in the environment^21^. To characterize the impact of these climatic factors, we used the Universal Thermal Climate Index (UTCI), which provides a one-dimensional measure of the sum of environmental heat stress and its effects on the human body^74^, as it can provide a more accurate description of the overall heat stress on biological organisms than just the individual temperature value. Data from the European Union’s Copernicus Earth Observation Programme shows that our southern study region experienced very strong heat stress during the vegetation period, while the North-Eastern control region only experienced strong heat stress in recent decades (**Fig. 16 A**)^75^. This distinction aligns with our findings of higher MT production in feedstuffs from the southern area (**Fig. 16 B)**. Specifically, levels of AF (p = 0.019), ZEA (p = 0.00051), and FB1 (p = 0.0018) were significantly elevated in the study area compared to the control region, though DON levels did not differ significantly (p = 0.12) (**Fig. 16 C)**. These data indicate a potential link between environmental heat stress and increased MT contamination in specific regions.

**Fig. 16.**
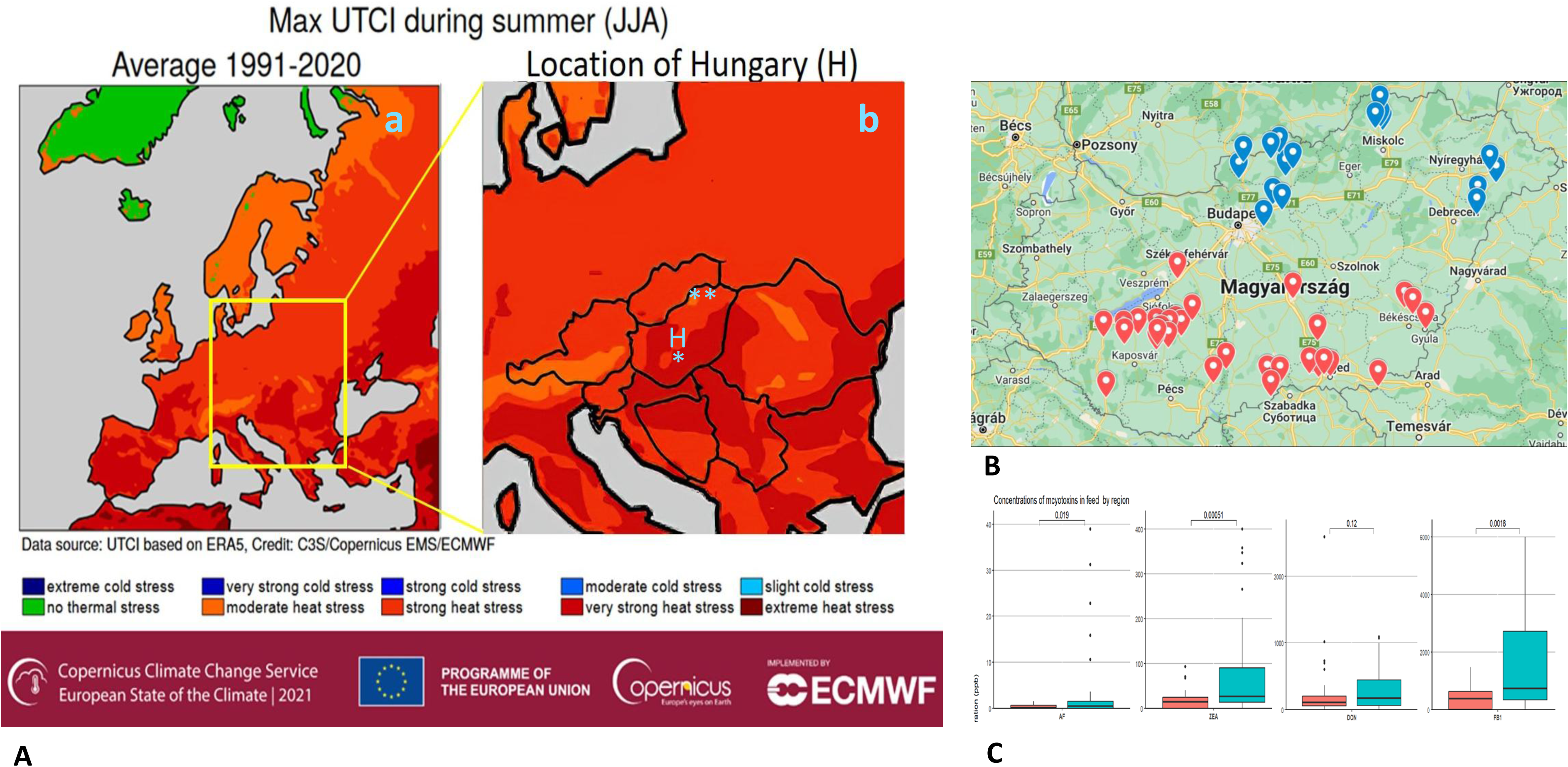
Geographical correlation between heat stress and mycotoxin load in Hungarian feeds. **A** The maximum Universal Thermal Climate Index (UTCI) measured in Hungary by European Union’s Copernicus Earth Observation Programme (using the https://climate.copernicus.eu/esotc/2021/heat-and-cold-stress). The UTCI, as a one-dimensional measure of the simultaneous effects of temperature, radiant heat, humidity and wind, was used by us, since we assumed it is the most accurate index of the thermal stress on biological organisms. **a** UTCI maximum averages summarized for the period 1991-2020 in Central Europe. **b** The borders of Hungary (H) and neighboring countries. The dark red color of the study southern area (asterisk) indicated very strong heat stress, but the red color of northeastern control region (duble asterisk) showed strong heat stress only. The color codes used by the Copernicus Earth Observation Programme. (Modified: https://climate.copernicus.eu/esotc/2021/heat-and-cold-stress.) **B** Sites of origin of feed samples from the Southern Hungary study region (red marks) and the control areas in Northern Hungary (blue marks) **C** The average mycotoxin values (aflatoxins AFs, zearalenone ZEA, deoxynivalenol DON, and fumonisin B1 FB1) of feed samples and their differencies on study and controll areas. AF (p<0.019), ZEA(p<0.00051), FB1(p<0.018) were significantly increased, but DON(p<0.12) was not. The area with very strong heat stress remarkable overlap with that has mycotoxin overproduction.

### Impact of Feeding Practices on Mycotoxin Exposure in Wildlife

Wildlife diet consists not only of natural vegetation but also supplemental feeding especially during the winter. It is especially popular among deer hunters in the South-Eastern US, practiced by 89% of Arkansas private land hunters ^76^. When animals rely on livestock feed, which can be contaminated by MTs either from field growth or suboptimal storage conditions ^19,77^. The test feed samples, representing both spreader and natural-sourced vegetation, were collected from October to February, spanning from harvest to winter’s end. Although estimating the exact intake ratio of these feeds is challenging, they are considered representative over the year. An observed practice involves scattering low-quality or moldy livestock feed for wildlife, often dispersed directly on the ground rather than placed in dedicated feeders. This approach significantly increases MT exposure ^77,78^. Due to the lack of standardisation possibility we did not include these lower-quality samples in our analysis, we hypothesize that such feed management practices contribute significantly to the high rates of pedicle and antler anomalies observed in wildlife populations.

### Antlers as Biomarkers

Antlers, renewed each year, provide a unique opportunity to assess accumulation of both organic and inorganic materials, making them valuable biomarkers for environmental monitoring, particularly in regions with dense cervid populations ^79,80^. The rapid annual division of pedicle stem cells, according to our work, can be highly sensitive to mycotoxins, so antlers can serve as indicators of environmental mycotoxin exposure. Therefore, investigating the abnormal growth patterns of antlers can provide an easily accessible method for assessing regional mycotoxin loads, reflecting environmental conditions and potential exposure risks ^15,17^.

Industrial pollutant emissions in Hungary have significantly decreased over the past two decades, particularly in the Southern regions^81^. Given this decline, we consider it unlikely that lead or fluoride exposure ^15,17^ is directly associated with the observed antler anomalies. This reduction in industrial pollutants suggests that other environmental factors may play a more significant role in the development of these anomalies.

### Limitations

Spanning a wide range of disciplines - meteorology, mycotoxicology, pathophysiology and pathoanatomy - our study looked at the impact of climate change on nature’s fastest regenerating bone tissue, the antler, which could not be covered in all its detail. We did not include examinations on the effects of MTs on cultured ASCs and could not carry out controlled animal feeding experiments supplemented with whole body autopsy. Not all of the RAPS characteristics describing PCD can be treated as independent variables and there can be conflicts between them, as discussed in the Methods. Resolving these requires consensus-based comparative studies on a larger number of cases. A correlation between mycotoxin production of fungi and the Universal Thermal Climate Index (UTCI), the best current descriptor of the effect of temperature change on human physiology, but its general applicability to other organisms requires further investigation.

### Summary

This study described a long-known but poorly characterised antler pedicle disorder, as a multimycotoxicosis-caused systemic disease, called Pedunculitis Chronica Deformans. Morphologically supporting Uwe Kierdorf and Horst Kierdorf’s hypothesis ^36,25^ which is based on Richard Goss’s early observation that scar and blastema formation are mutually exclusive^1^. However, when the former occurs this can lead to disease. The scar that forms in the casting wound inhibits antler regeneration, creating the potential for further complications such as pathological fracture or intracranial abscessation finally leading to animal death.

Our findings indicate that mycotoxins can be significant environmental contributors to this pathology, though establishing causality remains difficult due to various influencing factors, particularly the not properly evaluated health status of the affected animals. The study highlights the importance of accurately interpreting antler abnormalities and emphasizes the need for monitoring of environmental mycotoxin exposure as key strategies to mitigate this risk.

## Materials and Methods

### Institutional Review Board Statement

According to the statement of the Institutional Review Board (NAIK MBK MÁB 004-09/2018), the study is not considered as an experiment with animals because the researchers collected samples from the carcasses of legally harvested fallow deer hinds; consequently, the ethical treatment rules are not applicable. The carcasses were provided for the sampling by the authorized game managers, in full compliance with all ethical and legal regulations. Furthermore, Decision No. (TO-04H39/405-2/2020) of the Government Office of Tolna County, dated (05.06.2020), authorized the shooting of antlered fallow deer outside the hunting season for research purposes

### Source of anomalous and normal trophies

The study examined 50 *Dama dama* (DD) anomalous antler trophies collected between 2017 and 2021 from Somogy and Tolna counties in Southwest Hungary, Central Europe (study region), focusing on morphological deviations associated with Pedunculitis Chronica Deformans (PCD). A Morphology Control Group (MCG) served as the basis of comparison to this evaluation except for age, antler main beam length, and brow tine length. It comprised 24 trophies from Gúth (https://nyirerdo.hu/guthi-erdeszet/) where no antler anomalies were reported during the same period. This hunting area is located in Hajdú-Bihar County, in Northeastern Hungary, approximately 300 km northeast of our southern study region. Trophies in both groups ranged from 4 to 12 years in age. Additionally, five cases of roe deer (*Capreolus capreolus*, CC) and three cases of red deer (*Cervus elaphus*, CE) were analyzed.

The classic CIC (International Council for Game and Wildlife Conservation) trophy evaluation includes the assessment of age, antler beam length, and brow tine length measurements, therefore we could use these data from the Hungarian Trophy Register (HTR). From this, a Trophy Register Control Group (TRCG) was formed from 2,912 antler evaluation sheets, excluding those marked as “deformed,” “injured pedicle,” or “diseased pedicle”.

### Internal organs, blood and soft tissue samples

To establish a baseline for toxicological and hormonal parameters, blood, liver, and muscle samples were collected from 31 healthy DD during the hunting seasons of 2019 and 2020, serving as the control group. Additionally, samples were obtained from 58 animals exhibiting pronounced trophy abnormalities in the same period. The body condition of all animals was evaluated using a standardized four-point scale (poor, fair, good, excellent) according to the Hungarian Body Condition Index. Animals were further categorized into age groups: young, middle-aged, and old. For detailed morphological and histopathological examination, complete heads were collected from five DD individuals displaying aberrant antler morphology. Moreover, five DD ethically approved to be harvested in the velvet stage with pronounced antler anomalies were subjected to detailed anatomical analysis, including the antlers, surrounding skullcap, and associated soft tissues.

### Macroscopic examinations

A standardized protocol was developed for the macroscopic evaluation of trophies. An experienced wildlife manager (IL) with decades of trophy assessment experience and a consultant surgical pathologist (FS) conducted the initial assessments. To ensure inter-observer reliability, two additional trained observers (RK and AS) independently evaluated a subset of trophies. The morphological deviation of the trophy from the normal was evaluated for the roses (R), antlers (A), pedicle (P) and skull (S) with 3, 6, 7 and 6 parameters, respectively, for a total of 22 parameters on both sides in 24 healthy (MCG) and 50 abnormal cases **(Table 1.)**

**Table 1.** Detailed description of the RAPS features.

|  |  |  |
| --- | --- | --- |
| **Rose** | **R1** | Lack of continuity in the grain of the rose is longer than 5 mm but less than 1 cm. |
|  | **R2** | Rose deformity (also visible on cast antlers) |
|  | **R3** | Loss of rose granularity is greater than 1 cm |
| **Antler** | **A1** | Supernumerary tine originating from the rose, can be a sign of extra antler bud |
|  | **A2** | The length difference between the two sides of the brow tines |
|  | **A2.1** | Difference from Trophy Register Control Group (TRCG) which is derived from data of Hungarian Trophy Register after exclusions of cases marked abnormal |
|  | **A3** | The difference in length of the main beams relative to each other |
|  | **A3.1** | Difference relative to the TRCG |
|  | **A4** | Absence of antlers or aberrant antler beams. |
| **Pedicle** | **P1** | Pitting on the proximal surface of the pedicle |
|  | **P2** | The pedicle deformity characteristic to PCD in Fallow deer (*Dama dama*) is the deviation from the regular circle diameter more than 4 mm, also visible with the eye |
|  | **P3** | Shortening of the pedicle more than physiological |
|  | **P4** | Widening of the pedicle beyond normal extent |
|  | **P5** | Peduncle fistula |
|  | **P6** | Free casting plane from which does not protrude any antler-bony protuberance |
|  | **P7** | More than fifty percent of the casting plane of the pedicle is in or below the plane of the skull, considered to be the absence of it |
| **Skull** | **S1** | The pitting extends on the skull surface |
|  | **S2** | Spongy remodeling of skull bones where the cavities are larger than at intertrabecular spaces in spongy bone |
|  | **S3** | Discohesion along the interosseous suture |
|  | **S4** | Separation of the cortical bones independently from a sign of bone displacement |
|  | **S5** | A fistula mouth opens distant from the pedicle on the skull |
|  | **S6** | The center of the pedicle has shifted significantly, mostly towards the brow ridge |

### RAPS features

Furthermore, the RAPS features were assorted into 4 severity grades based on their pathomorphological background **(Table 2)**

**Table 2.** Detailed description of the RAPS features.

|  |  |
| --- | --- |
| **Grade I** | R1, A1, P1, S1 |
| **Grade II** | P2, R2, A2, A2.1, P2, P3, P4, S2, S6 |
| **Grade III** | R3, A3, A3.1, P5, P6, S4 |
| **Grade IV** | A4, P7, S3, S4 |

### Considerations for RAPS score assessment

Despite their rigorous applicability, the RAPS features cannot be treated as entirely independent variables due to inherent conflicts and contingencies. For instance, R1 and R2 are mutually exclusive events. Additionally, A2 and A3 were determined by comparing the two sides, and to maintain consistency, we assigned a score of 1 on both sides if present. However, this approach introduces an inherent bias into the downstream analysis by affecting the evaluation of laterality. Furthermore, our observations indicated that the age-stratified brow tine and main beam length distributions in the Trophy Register Control Group (TRCG) were often skewed to the left. As a result, for the assessment of A2.1, we implemented a less stringent bottom 20% threshold, which allowed for a greater than 15% increase in sensitivity for this marker.

### Analysis of Pedicle Ellipticity

We tested whether PCD induces elliptical distortion of the pedicle using the minor (d) and major (D) axes of pedicles (modelled as ellipses). Ellipticity was quantified using the distance of foci from the origin [inline] and eccentricity (*foci distance/D*).

Both metrics revealed significantly different means between the sick and healthy cohorts (2 sample Wilcoxon tests, p-values: 4.371*10^-7^ and 1.426*10**^-4^**respectively.)

### Survival Analysis

The Kaplan-Meier survival analysis was conducted on the PCD, MCG, and TRCG cohorts, with the log-rank test used to assess the statistical significance of survival differences across these groups. The analysis revealed that the sick cohort exhibited significantly poorer survival compared to both the TRCG and MCG cohorts.

### Microscopic Examinations

5 DD heads with soft parts (skin, connective tissue, muscle and brain) were collected in the velvet phase and with dry antlers each, showing obvious abnormalities at least one side. The proximal ca. 20 cm long segment of the antler, the whole antler in case of obvious abnormality and pedicle attached to the antler and the surrounding skull roof, including all soft tissues and brain were cut out and fixed in 10% formalin for at least one week. These preserves were sawed into 4 mm thick slices with a meat industry band saw. Visible alterations were excised for microscopic examination. Electric decalcification using a Tissue-tek TDE 30 Decalcifier system (ref.:1428 Sakura Finetek Europe, Alphen aan den Rijn, Netherlands,) was used for three weeks with continuous monitoring. Samples that couldn’t be sectioned after this time were further handled for one month in a solution of 70% ethanol containing 5% phenol or Q path DC3 (VWR ref.:09128300. Vienna, Austria) for three weeks.

After decalcification, traditional pathological techniques (embedding in a paraffin block and making 5 micrometre sections) were employed, including Hematoxylin & Eosin, PAS, Warthin-Starry, Giemsa, Zhiel-Neelsen, and Wade-Fite staining to detect potential pathogens. In four cases of one year-old trophies, parallel 4 mm thick slices were sawed from the pedicle and after decalcification, microscopic examinations were conducted.

### Radiological examinations

Computed Tomography (CT) images of 16 deer trophies and 2 complete heads with soft tissue were acquired with a single slice GE CT Scanner Model CT/e. Acquisition parameters for CT images were as follows: helical mode, 512 × 512 matrix, 1 mm slice thickness, 1 mm image interval, 120 kVp, 20-120 mA, and bone convolution kernel. The preparations were placed in a prone position on the CT bed with the aid of foam wedges. Transverse images of the skull were obtained perpendicular to the hard palate. The scanned area ranged from the occipital bone to the mid-orbital region including the antler tree. The osseous structures on DICOM images were evaluated using a window width of 4000 and level of 400 HU.

### Microbiological Investigations

From 2018 to 2020, we conducted microbiological examinations on 42 antler pedicle problem cases of fallow deers (*Dama dama*). These investigations included both traditional culturing methods and next-generation sequencing (NGS) to characterize the bacterial genome and describe the proportions of pathogens present.

The swabs collected from inflammatory exudate samples were cultured on 5% sheep blood agar under both aerobic and anaerobic conditions at 37°C for up to 5 days. Direct smear examination using Gram staining was performed on all samples from infected tissues. Bacterial isolates were identified based on their Gram stain reaction, cultural properties, morphological characteristics and results from standard biochemical tests.

Prior to metagenomic approaches, inflammatory exudate-covered peripeduncular soft tissue homogenates were exposed to nucleic acid extraction with Quick-DNA Fungal/Bacterial Microprep Kit (Zymo research). Libraries were prepared using the PCR Barcoding Kit (SQK-PBK004) and protocol by Oxford NanoporeTechnologies. The sequencing was performed on the R9.4.1 flow cell with the usage of MinKNOW 19.12.5 software. The basecalling was carried out with GUPPY in real-time with the usage of fast base calling algorithm (used config file: dna_r9.4.1_450bps_fast.cfg). After the sequencing the data were demultiplexed with Guppy barcoder and adapters were trimmed using Porechop v0.2.4 [a] with default settings. Internal adapters were not allowed because they have a strong indication of chimeric reads. These reads were not used in the further analysis. All remaining reads were aligned against the full NCBI NR (National Center for Biotechnology Information non-redundant protein) database with the usage of DIAMOND software which is suitable for long reads^82^. Bacterial sequences were extracted and grouped into levels of genus from the DIAMOND results.

### Analysis of Toxicological and Hormonal Parameters

#### Mycotoxin Analysis of Serum and Tissue Samples

Mycotoxin (MT) levels in serum and tissue samples (liver, muscle) were measured by ELISA, the optimization of which has been described previously^83^. FB1, ZEA, DON, and total AF (B1, B2, G1, G2) were assessed through immunoassays.

Total AFs and DON concentrations were measured using Toxi Watch ELISA kits (Soft Flow Ltd., Pécs Hungary), validated by the manufacturer for serum samples and various organs and tissues ^51,84,85^. Serum samples were thawed, diluted, and extracted with a threefold EtOH/water solution (23/77, v/v) shaken for 15 minutes at room temperature. Supernatants were collected, and DON was diluted 10 times with 0.01 M PBS, pH 7.4. All samples were measured in triplicates. ZEA was analyzed using Ridascreen Zearalenone Enzyme Immunoassay kits from R Biopharm (Art No.: R1401, Arnhem, Germany). Serum samples were prepared using the RIDA© C18 Column (Art No.: R2002, R-Biopharm, Arnhem, Germany) per the manufacturer’s instructions and measured in triplicate. Liver tissue was homogenized with a FastPrep-24 Classic homogenizer (MP Biomedicals, Irvine, CA, USA) in ice-cold 50 mM sodium acetate buffer (pH 4.8), incubated for 3 hours at 37°C with Helix pomatia β-glucuronidase/arylsulfatase (BGALA-RO, Roche, Basel, Switzerland) per the manufacturer’s instructions. Extraction used 70% methanol and 30% water (v/v), and extracts were centrifuged at 8000xg for 5 minutes at room temperature. The supernatants were diluted with an assay buffer. Measurements were taken using a Thermo MultiskanTM FC microplate reader (Waltham, MA, USA) with SkanIt RE software (version 6.1.1.7) at 450 nm absorbance with a 630 nm reference wavelength. FB1 was quantified using the EUROPROXIMA Fumonisin (5121FUM) assay kit (R-Biopharm, Arnhem, Germany), validated for serum and various animal tissues. The manufacturer’s instructions were followed, and triplicate measurements were performed for each sample.

#### Mycotoxin Analysis of Feed Samples

The test feed samples came partly from the spreader and from the natural vegetation of the given area. The collection period spanned from October to February, from harvesting to the end of winter. The mycotoxin content in the feed samples was quantitatively determined using the Fungi-Plex™ multiplex mycotoxin assay kit (Soft Flow, Ltd., Pécs, Hungary), which employs microbeads and flow cytometry for the simultaneous qualitative and quantitative detection of six mycotoxins: aflatoxin B1 (AB1), zearalenone (ZEN), deoxynivalenol (DON), fumonisin B1 (FB1). Details of the Fungi-Plex™ measurements were described by Czéh et al.^83^ using the FACSArray™ BD Bioanalyzer FC with onboard acquisition software, both from BD Biosciences (Erembodegem, Belgium). MultiScreen HTS-BV 1.2 μm clear non-sterile disposable filter plates were utilized for all experiments, in conjunction with a Heidolph Titramax 101 platform shaker and a MultiScreen HTS Vacuum Manifold, both from Merck (Budapest, Hungary). The additional onboard software used was FCAP Array v3.0.^86^

#### Hormone analyses

Hormone analyses followed the manufacturer’s guidelines. Serum samples were collected during rutting seasons, measured in triplicate using the 17-beta-estradiol kit (Cat No: DNOV003), progesterone kit (Cat No: DNOV006), and testosterone kit (Cat No: DNOV002), all from NovaTec Immundiagnostica, Dietzenbach, Germany

## Statistical Analysis

The presence of morphological anomalies (R, A, P, S) was recorded with a value of 1 for presence and 0 for absence, separately for each side. A2.1 and A3.1 were considered as present if the brow tine and main beam length were lower than the bottom 20% and 5% quantiles of the distributions of brow tine and main beam length in the TRCG of the corresponding age and side. Pairwise comparisons between means unless stated otherwise, were carried out using 2-sample Wilcoxon tests, with Holm’s multiple testing correction. Computations were performed using the R language and environment^87^, figures were drawn using ggplot2 ^88^, ggpubr ^89^, ggraph^90^ and pheatmap^91^.

## Authors contribution

F.S., I.L., L.S., and Z.S. conceived and designed the study. F.S., A.U., T.P., and R.K. prepared the original draft, while S.F., B.N.B., P.G., Mi.Me., K.P., A.S., and F.K. contributed to review and editing. I.L., G.P., S.D., and Mi.Mi. managed the collection of anomalous and normal trophies. F.S., I.L., R.K., and A.S. performed the macroscopical assessments. F.S. and R.K. conducted microscopic examinations. Z.S., B.B., G.S., P.P., Z.M., and A.C. conducted toxicological investigations of internal organs, blood, and soft tissue samples, while B.B. and G.S. performed hormonal parameter analyses. A.C. and G.N. collected and analyzed feed samples. A.A.T. and K.E. performed radiological assessments, and A.A.T. and K.E. also carried out microbiological investigations, with G.K. conducting metagenomics analysis. A.U., G.H., A.H., D.T., and Ma.Ma. performed statistical analyses. L.S. and Z.S. supervised the study, provided critical revisions, and secured funding. All authors contributed to data interpretation and approved the final version of the manuscript.

## Acknowledgments

The authors of this article extend their appreciation to Gyulaj Forestry & Hunting Plc. for their support.

The authors are grateful for Péter Zámbó, the State Secretary for Forestry and Land Affairs. The authors also thank Sándor Juhos, Bence Fábián for their contributions to photography.

## Figure legends

**Supplementary Fig. 1.**
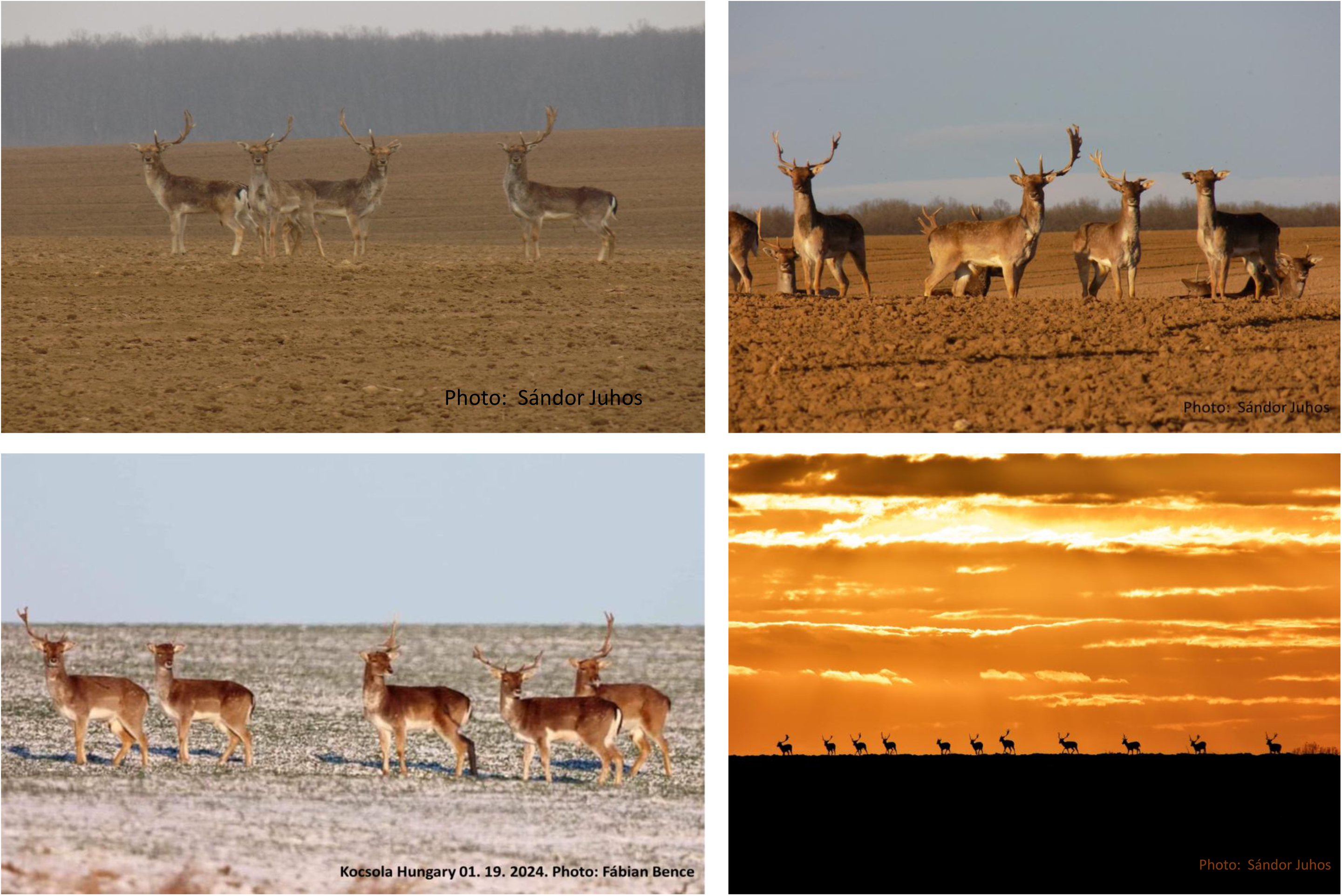

**Supplementary Fig. 2.**
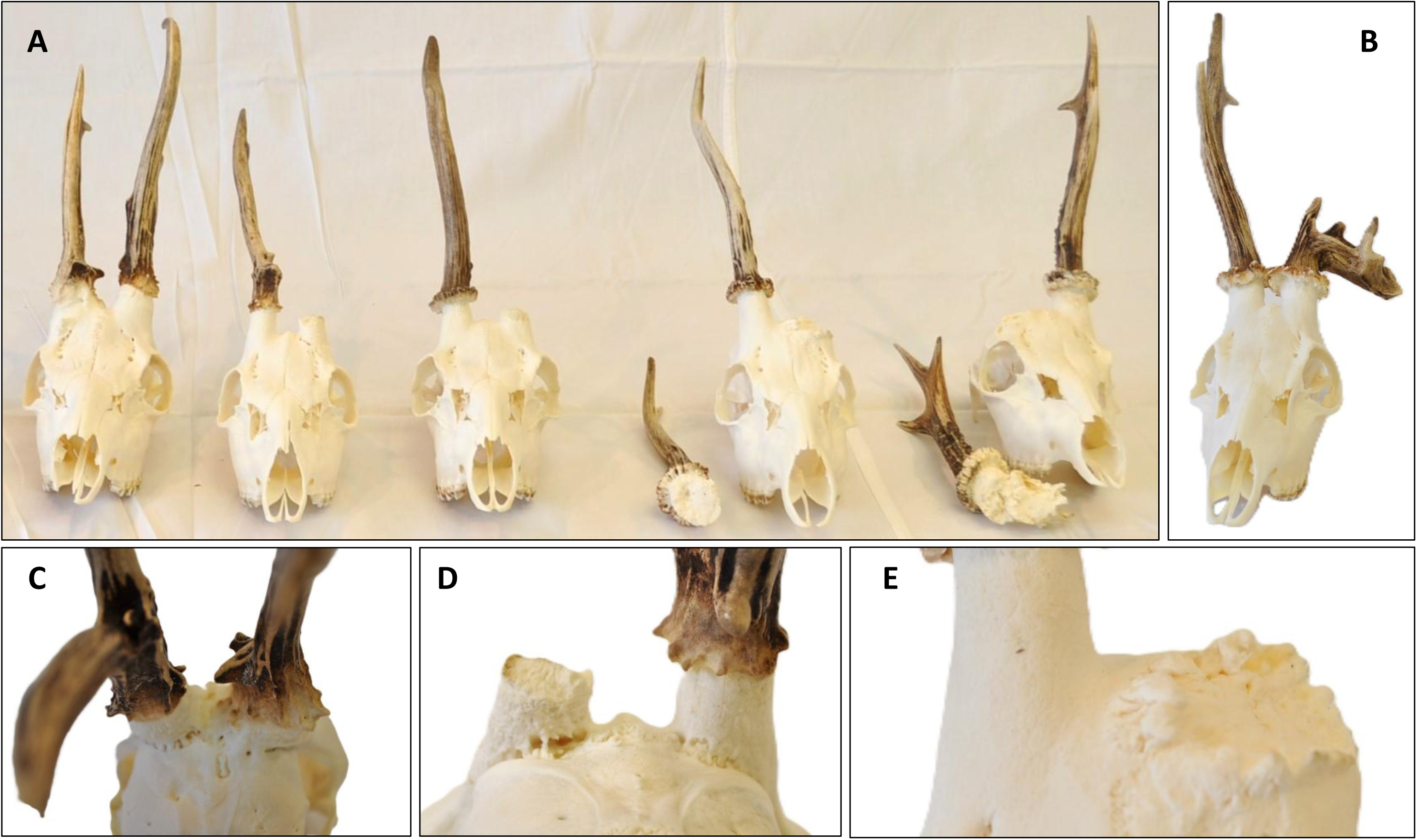
Pedunculitis Chronica Deformans (PCD)-caused trophy anomalies on roe deer (*Capreolus careolus*, CC). **A**: Our five cases put in order of severity show some signs of PCD. **B**. One velvet phase injured CC without visible rose, pedicle or skull alterations. **C**. The CC case 1 seen from behind. There are no brow tines nor antler burr on both sides. **D**. The CC case 2 seen from behind. The distal pedicle surface shows pitting on both sides, more prominently on the side from where premature casting happened. On this side note teh fissura formation between the pedicle and frontal bone, and on the other side the loss of burr granulation. **E**. On the left side of CC case 4 a margin crest developed on the broadening pedicle base, showing the osteoblastic bone building activity. The effect of osteoclasts does not step through the pedicle opposite what we can see by roe deer (*Capreolus careolus*, CC) or, even more prominently, by red deer (*Cervus elaphus*, CE).

**Supplementary Fig. 3.**
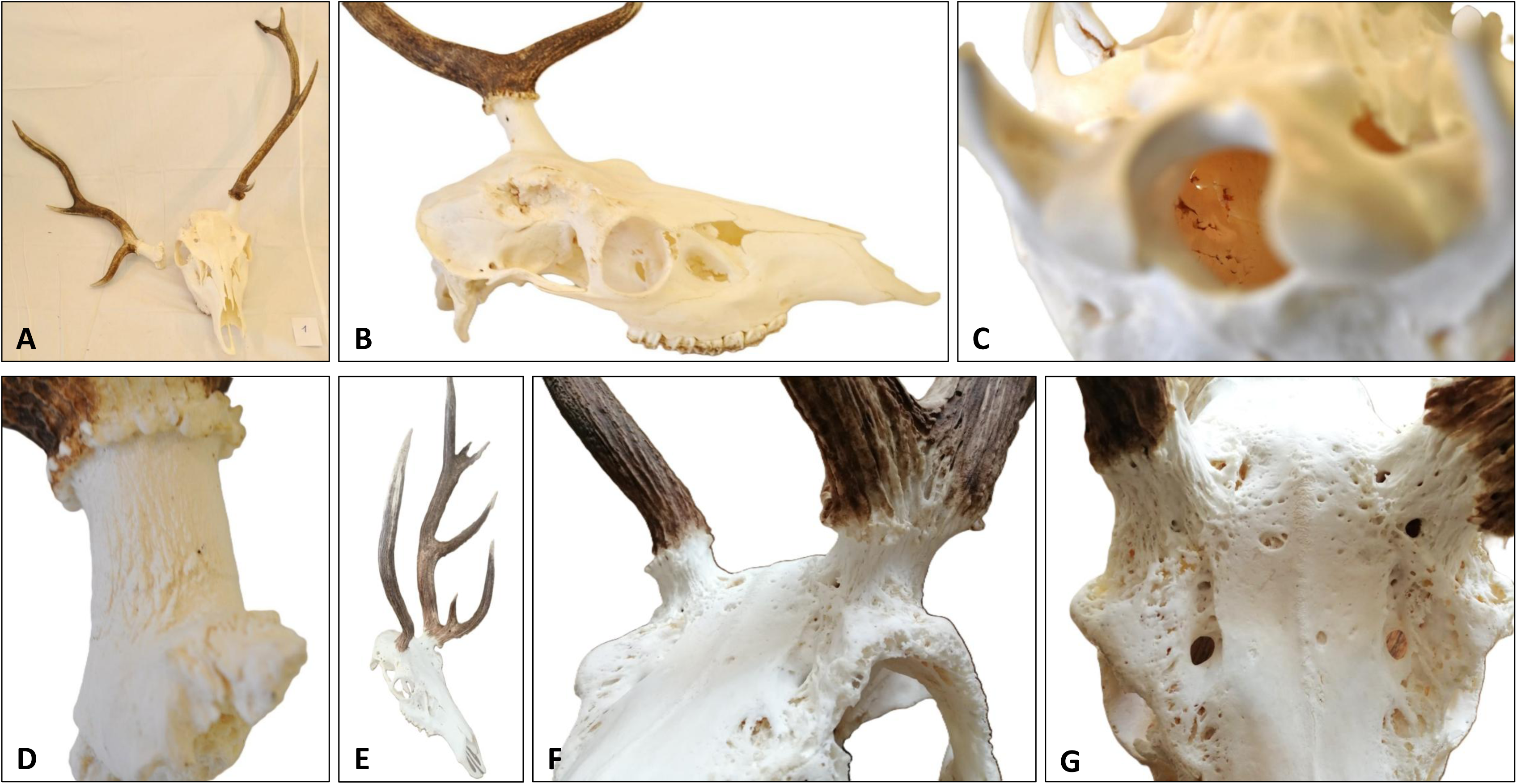
Signs of Pedunculitis Chronica Deformans (PCD) appearing on red deer (Cervus elaphus, CE). **A-D**: A two year old stag trophy with a broken out right pedicle serves as an example of early phase PCD where aseptic bone necrosis lead to pathological fracture. The right mean beam is shorter and less developed than the left one. The break happened during the hunt therefore we could examine both sides. **B**. The broad breaking surface involved most of the frontal bone, approaching the orbital arch too. **C**. View through the *foramen magnum* towards the right pedicle base. The spongiform hole formation happened without displacement of the skull bone surface and was not situated along a line which could be a sign of a traumatic impact. **D**. The broken out pedicle shoved remarkable pitting on the proximal two-thirds of the pedicle surface without its deformation or loss of rose granulation. **E-G** A case of the five years old stag as an example where the inhibited cast wound healing could lead to PCD. **E**. The difference between two sides is prominent, on the right, as only an aberrant antler beam developed. (There is no brow tine, therefore this structure cannot be considered as an antler.) **F**. Total loss of rose granularity on both sides without pedicle shortening/broadening/deformation but surface irregularity is obvious. **G**. The spongious remodelling due to osteoclast activation is a hallmark of PCD in CE. On both sides, more of the frontal bones are involved and the disease is reaching the orbital arches too. Opposite to fallow deer (Dama dama) there is no bony crest formation and broadening of the pedicle, which could be the signs of elevated osteoblast activity.

**Supplementary Fig. 4.**
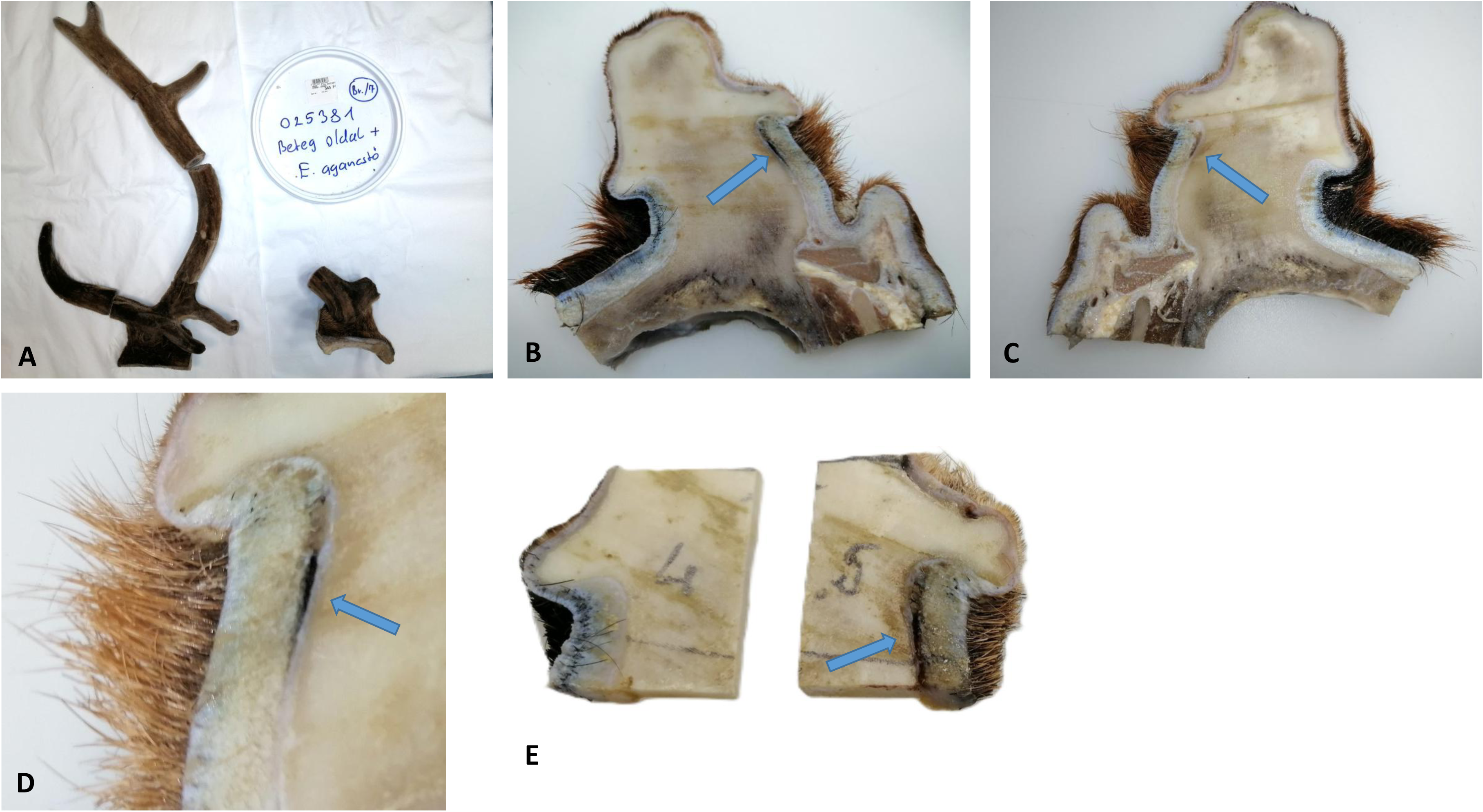
Example of Rose, Antler, Pedicle, Skull (RAPS) scoring. 1: Symptom present, 0: Symptom absent (Only anomalie present on the speciment are marked) **a** Posterior view of right rose and pedicle. **R2:** Rose deformity. **R3** Rose Granularity absence > 1cm. **b** Anterior view of right pedicle and skull surface. Pitting visible on pedicle (P1) and on the skull (S1) surfaces. **c** Trophy of a nine year old fallow deer (Dama dama) stag with remarkable asymmetry. According to our definition of antlers (main beam is longer than the brow tine, both originated from own buds and at least a rudimentary rose) the right side is not an antler but an Aberrant Antlered Beam (AAB), because the frontal bud has not developed into a frontal branch. Next to this is a supernumerary tine (A1) which grows from the medial (not anterior as brow tine) area of the rose. This can be a sign of an extra antlerogenic bud. Therefore, by comparing the brow tines of the two sides to each other (A2) and to the Trophy Register Control Group (A2.1), both of these symptoms can be socred as present on this specimen. The difference in length between the right AAB and the left main beam is significant by visual comparison (A3). **d** Right pedicle from medial side. Pedicle fistula with metal probe (P5) where the fistula mouth opens distant from the pedicle on the skull (S4). **e** Right pedicle from lateral side. Spongious remodelling (S2) with skull cortical bone opening (S5) and pedicle dislocation to orbital arch (S6) is visible.

**Supplementary Fig. 5.**
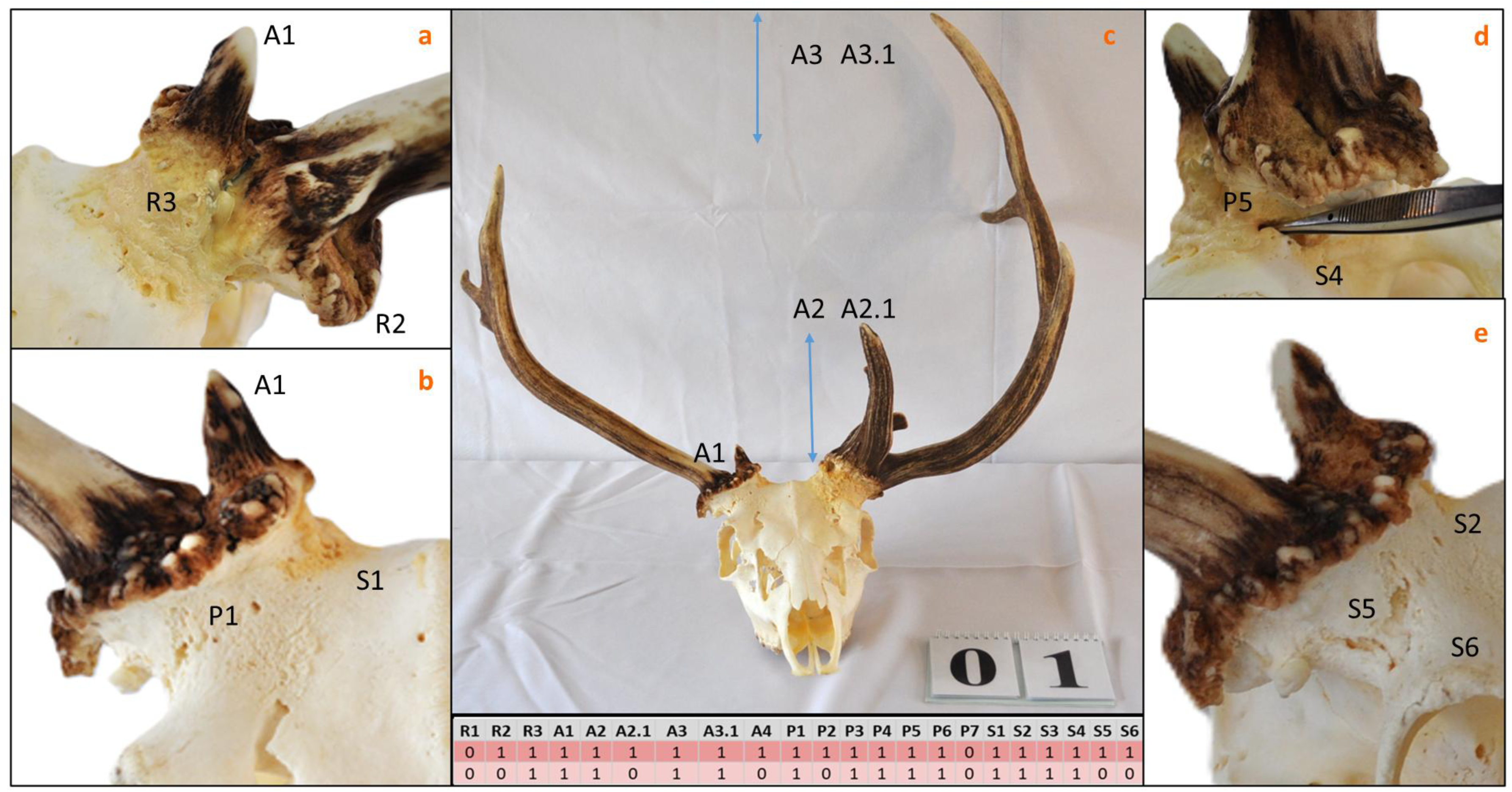
Fallow deer (*Dama dama*) in velvet. Antler abnormalities are visible at all stages of antler development. **A** At the earliest stage, a three-way divided antler forms a with a rudimentary brow tine and a retarded main beam on the right side. **B** On the right side an Aberrant Antler Beam (AAB) developed only. The main beam is substantially shorter than the brow tine, therefore we do not consider this appendage as an antler (A4). **C** On the right side, a pedicle deformity (P2) is obvious and a supernumerary tine (A1) is also visible. A longer brow tine (A2) can be a sign of compensatory growth (see text). On the left side an AAB is visible only without rose formation. **D** On the left side a prominent supernumerary tine (A1) is visible with a short brow tine (A2) and main beam (A3), where the latter is longer than the former, therefore we consider it as an antler**. E** On the left side a supernumerary tine (A1) exists which is shorter than the main beam on the other side (A3). **F** On the right side next to the supernumerary tine (A1) the main beam shortening (A3) is conspicuous**. G** An AAB with a fistula opening on the eyebrow arch (S4) (asterisk).

**Supplementary Fig. 6.**
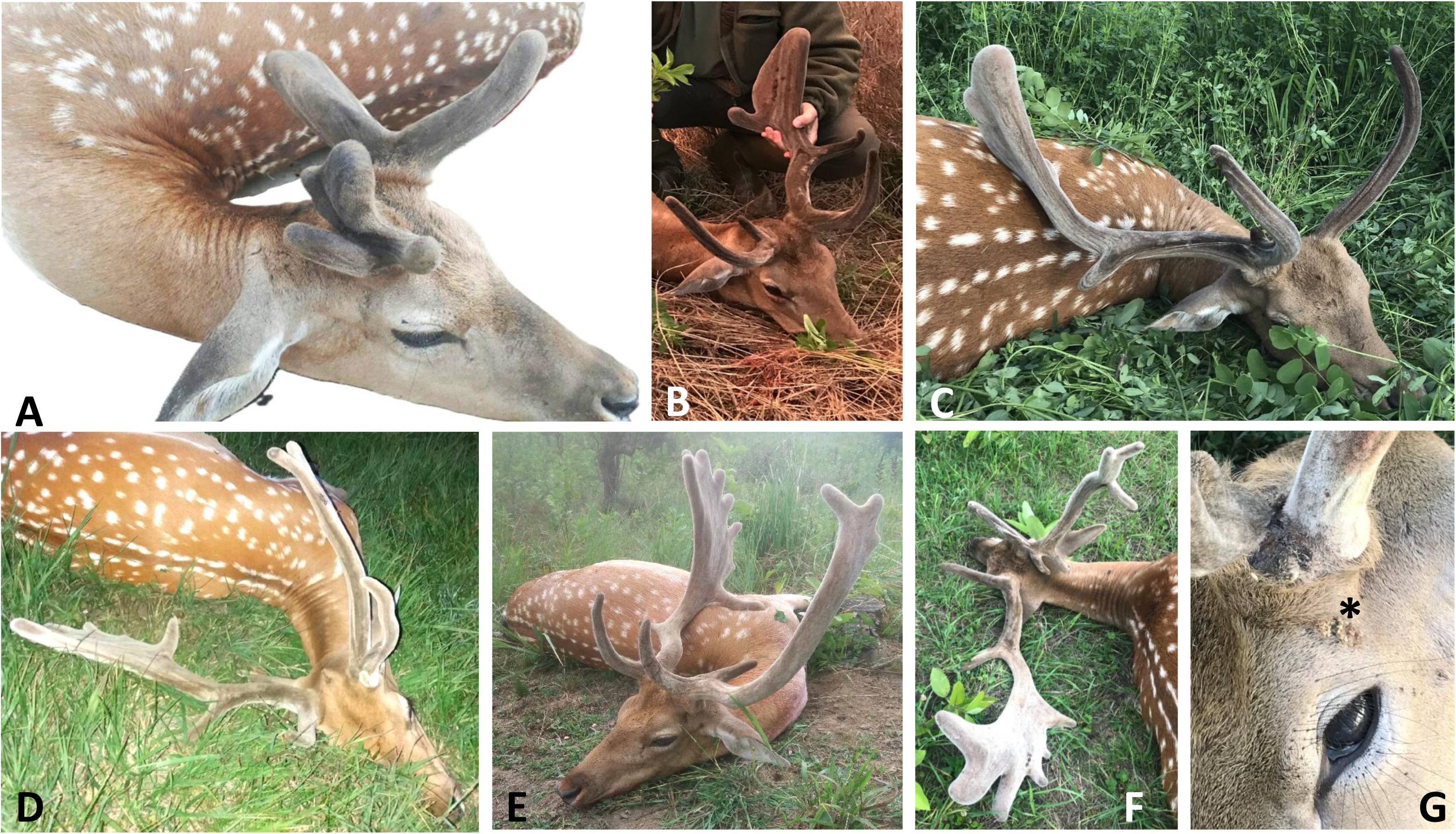
Manifestation of Pedunculitis Chronica Deformans (PCD) on harvested fallow deer (*Dama dama,* DD). **A** The antler-like protrusion on the left side may give the appearance of a post-breakout condition. However, the deformed rose (R2) with lack of granulation (R3) could only be the result of disease. After the dry antler is broken off, the remaining dry (dead) antler parts can no longer change into an abnormal shape. Hyperpigmentation of the skin of the right pedicle may indicate underlying inflammation, which may be a sign of a pedicle pitting (P1). **B** Next to the supernumerary tine (A1) the rose deformity (R2) and loss of rose granularity greater than 1 cm (R3) are obvious. Our study shows that R2 and R3 can also be informative for subcutaneous lesions. Loss of hair around the pedicle and purulent exudate are signs of superinfection. Next to the rose anomalies, R2 and R3 and pedicle P2 (deformity), P3 (shortening), P4 (widening) aberrations appeared on the right side with pedicle dislocation on the skull towards the eyebrow arch (S6). On the other side, A1 appeared, which can be a discrete sign of compensatory growth. **C** On the right side an Aberrant Antler Beam (AAB) is visible (there is no brow tine) with R2 and R3 and obvious positioning of the pedicle towards the eye brow arch (S6). On the other side A1 is also apparent. **D** On the left side AABs are visible with R3, P2, P3 and P4 malformations. Despite the presence of an antler palm we do not consider it as an antler due to the lack of a brow tine. **E** After shooting the three-year-old DD buck, the right pedicle broke out. This would not normally happen, suggesting a predisposing disease, as confirmed by the uneven fracture surface of the pedicle on the level of the skull. **F** We considered the right side as an AAB only, regardless of the presence of antler palm since the brow tine is lost. On the other side visible protrusions can only be classed as an AAB. **G** Same situation as C but the skin hyperpigmentation here is more prominent supporting the suggestion asserted with case A.

**Supplementary Fig 7.**
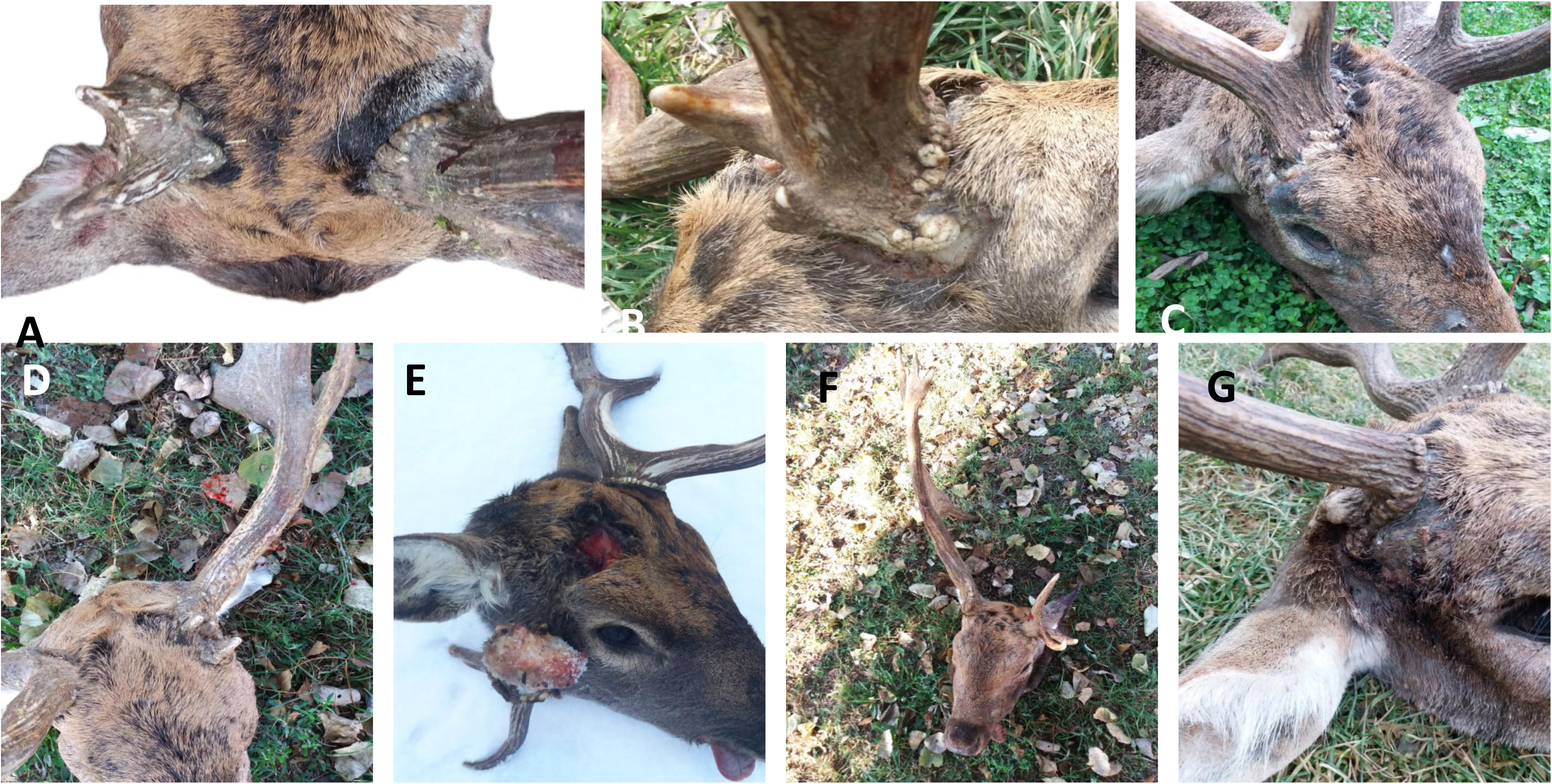
Manifestation of the Pedunculitis Chronica Deformans (PCD) on harvested Fallow deer (*Dama dama,* DD). **A** The antler-like protrusion on the left side may give the appearance of a post-breakdown condition. However, the deformed rose (R2) with lack of granulation (R3) could only be the result of a malformation. After the dry antler is broken off, the remaining dry (dead) antler parts can no longer change into an abnormal shape. Hyperpigmentation of the skin of the right pedicle may indicate underlying inflammation, which may be a sign of a pedicle pitting (P1). **B** Next to the supernumerary tine (A1) the rose deformity (R2) and loss of rose granularity greater than 1 cm (R3) are obvious. Our study shows that the R2 and R3 can also be informative for subcutaneous lesions. Loss of hair around the pedicle and purulent exudate are the signs of superinfection of PCD. Next to the rose anomalies, R2 and R3 and peduncle P2 (deformity), P3 (shortening), P4 (widening) aberration appeared on the right side with pedicle dislocation on the skull to the eyebrow arch (S6). On the other side a A1 appeared, which can be a discrete sign of compensatory growth **C** On the right side AAB is visible (there is no brow tine) with R2 and R3 an obvious positioning of the pedicle towards the eye brow arch (S6). On the other side a A1 is formed. **D** On the left side Aberrant Antler Beams (AAB) are visible with R3 and P2, P3,P4 malformations. Despite the presence of antler palm we do not consider it as an antler since the lack of brow tine. **E** After shooting the three-year-old DD buck, the right pedicle was broken-out. This would not normally happen, suggesting a predisposing disease, as confirmed by the uneven fracture surface of the pedicle on the level of the skull. **F** We consider the right side as an AAB only, regardless of presence of antler palm since the brow tine is lost. On the other side visible protrusions can be interpreted as an AABG Same situation as C but the skin hyperpigmentation here more prominent supporting the suggestion what is visible on case A.

**Supplementary. Fig. 8.**
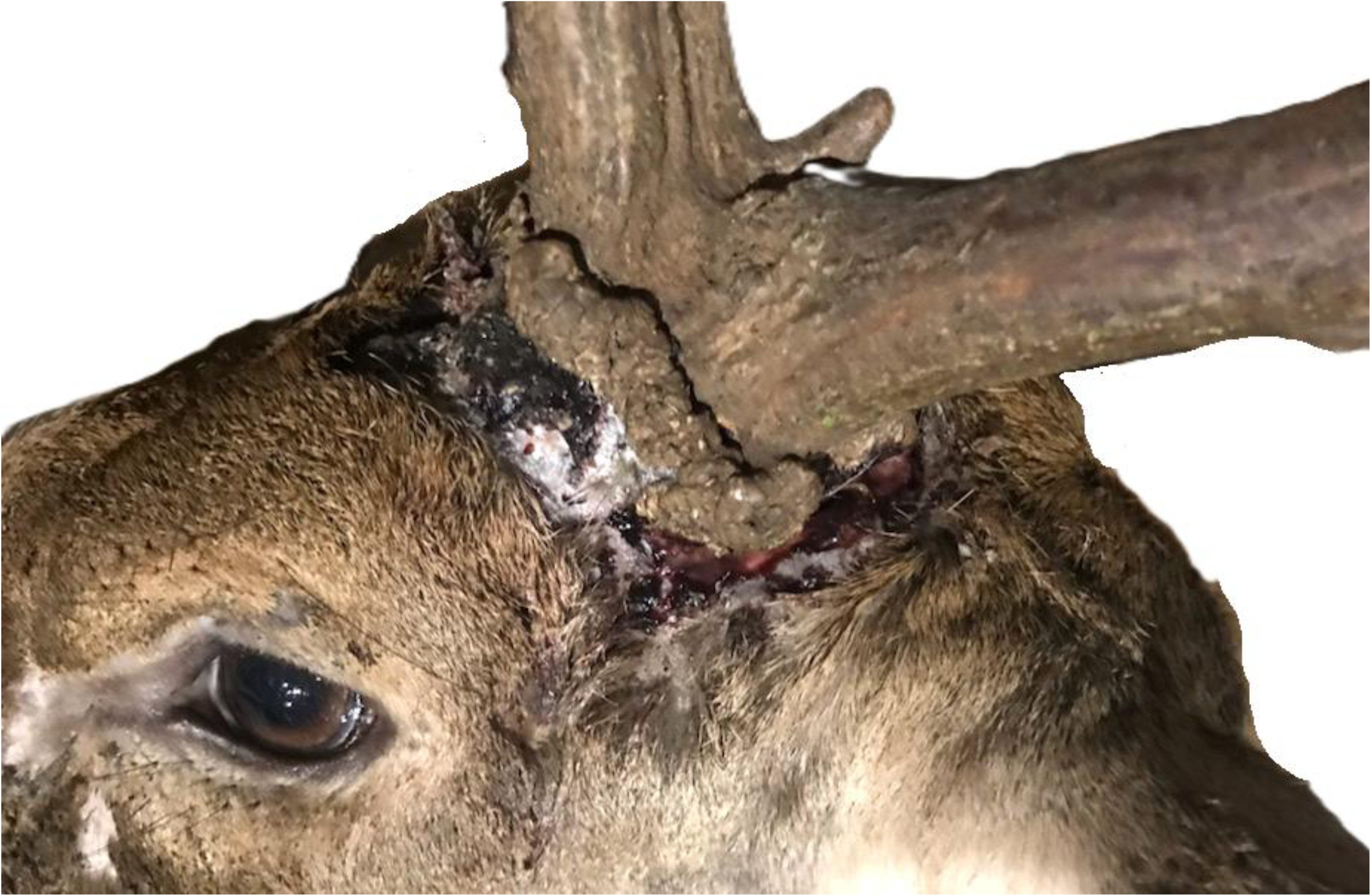
A phenomenon known in hunting communities as „pedicle rot”. The skin of the antler is covered with a hemorrhagic-purulent exudate, a sign of superinfection with suppurative bacteria, which is not a cause but a consequence of the underlying disease. The examination of RAPS (Rose (R), Antler(A), Pedicle(P), Skull (S) features help the recognition of Pedunculitis Chronica Deforman (PCD): Note the rose deformity (R2), granularity discontinuity (R3), the supernumerary tine (A1) the deformity (P2) shortening (P3) and widening (P4) of pedicle. A slight dislocation of the pedicle on the skull (S6) can be assumed.

**Supplementary Video. 1.**
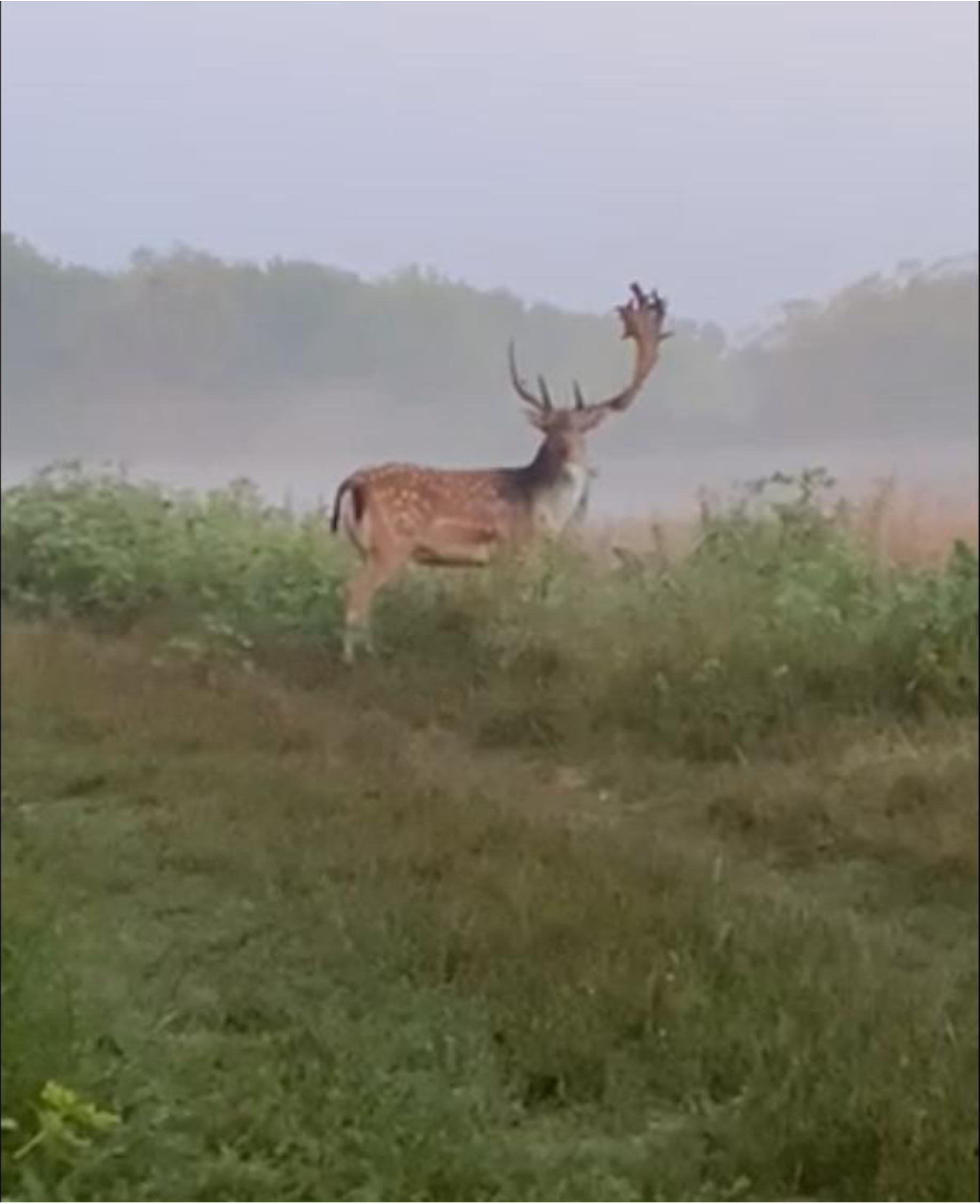

**Supplementary Data 1.**
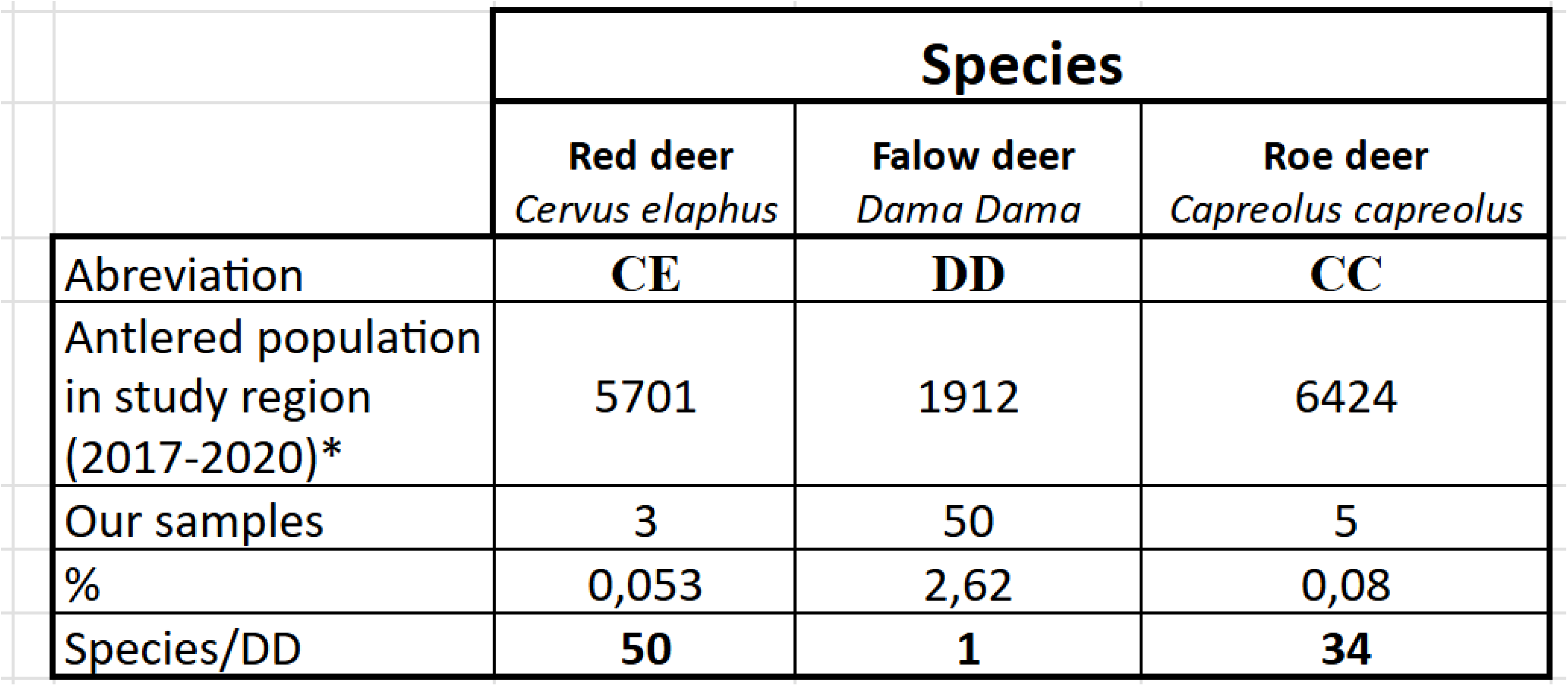

**Supplementary Data 2.**
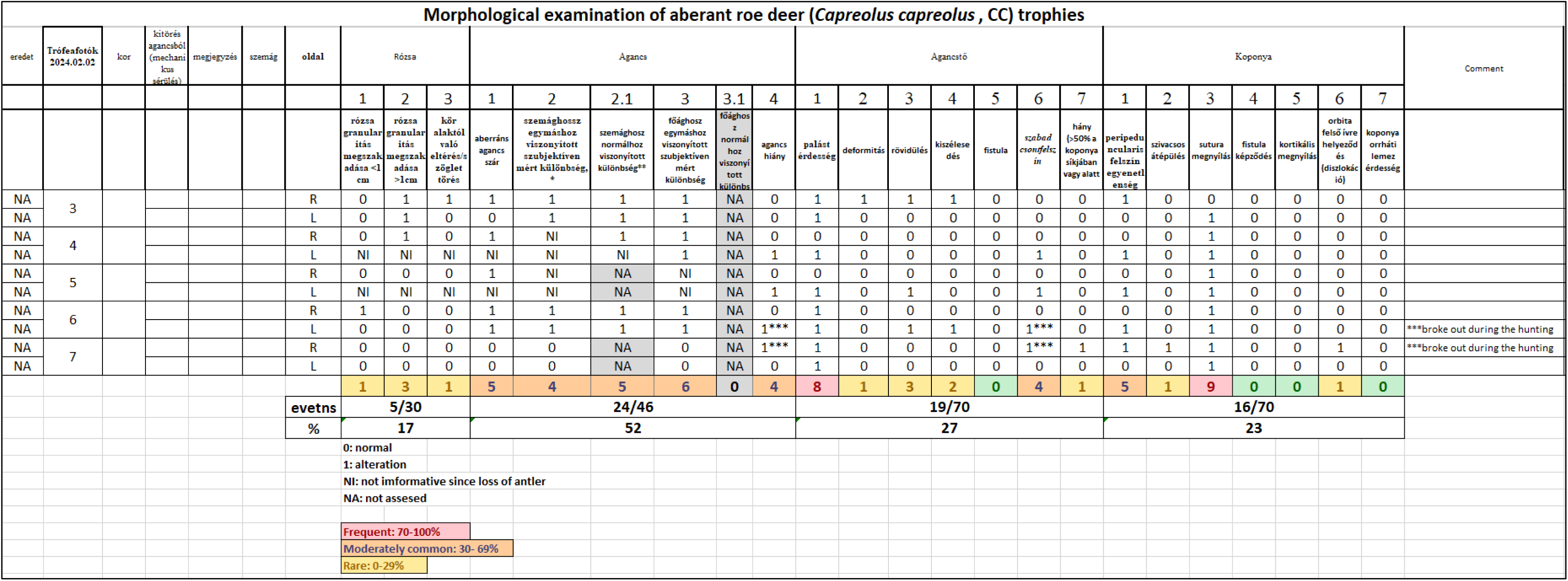

**Supplementary Data 3.**
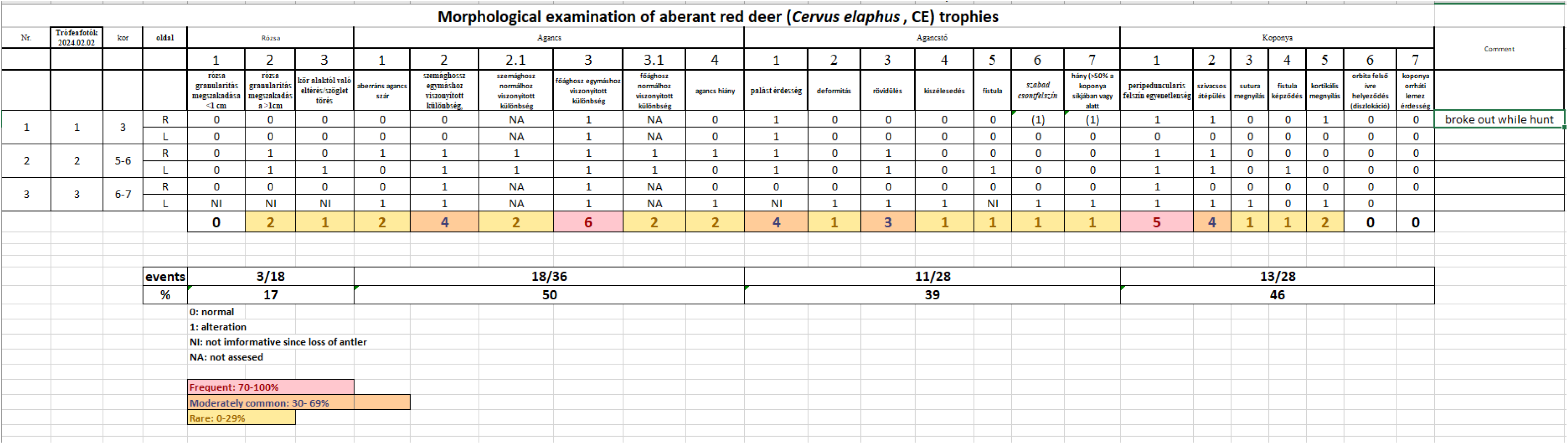

**Supplementary Data 6.**
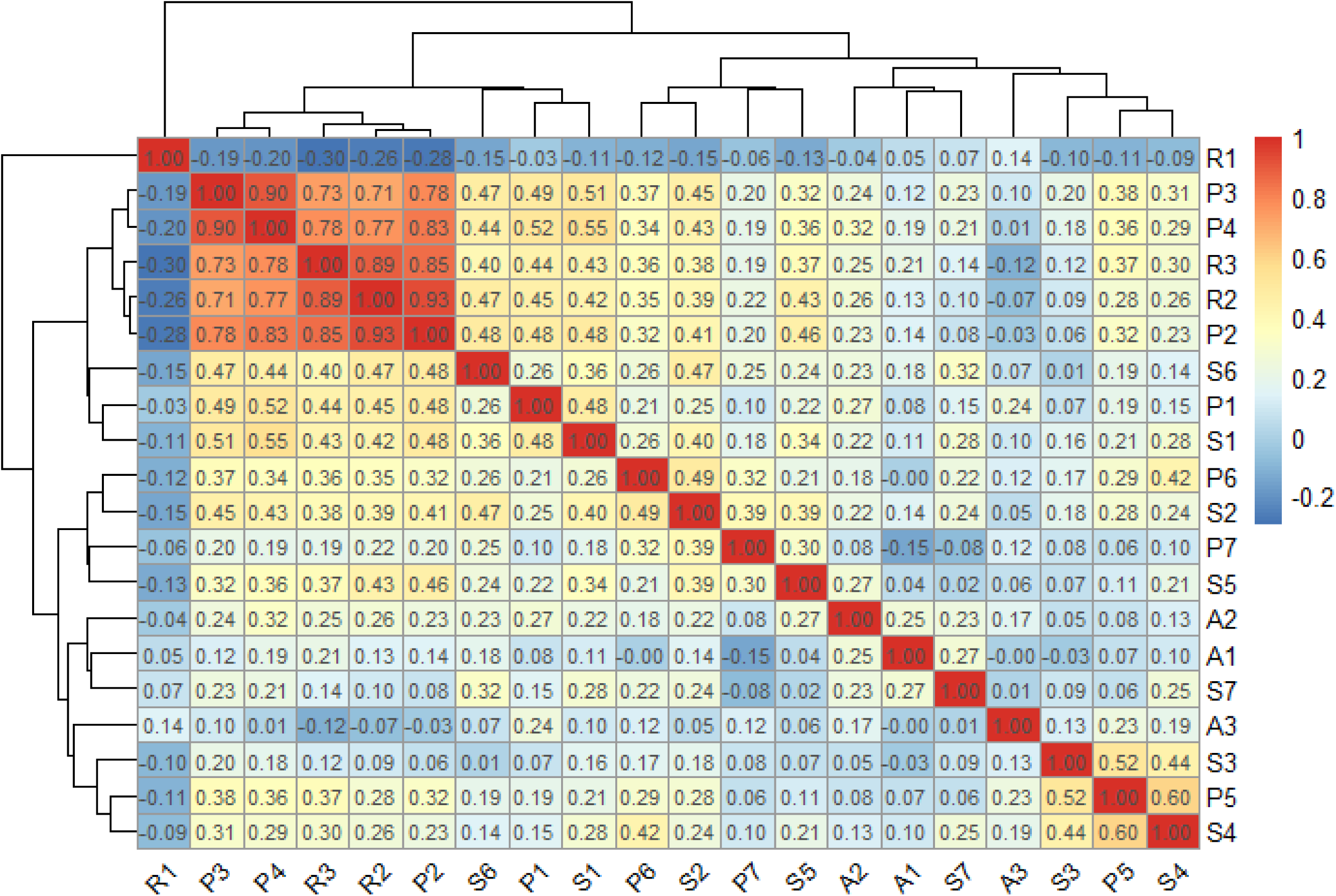

**Supplementary Data 7.**
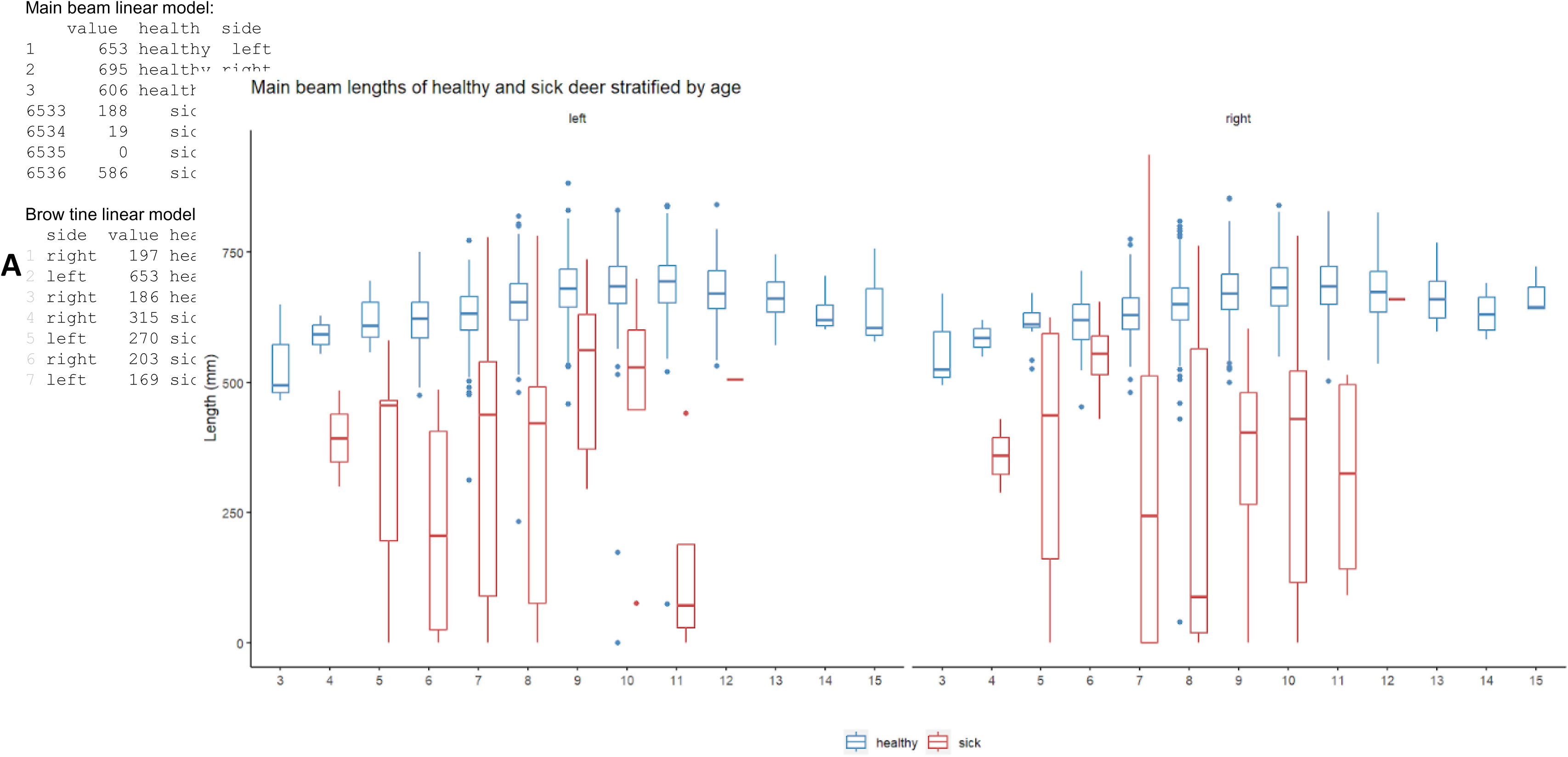

**Supplementary Data 9.**
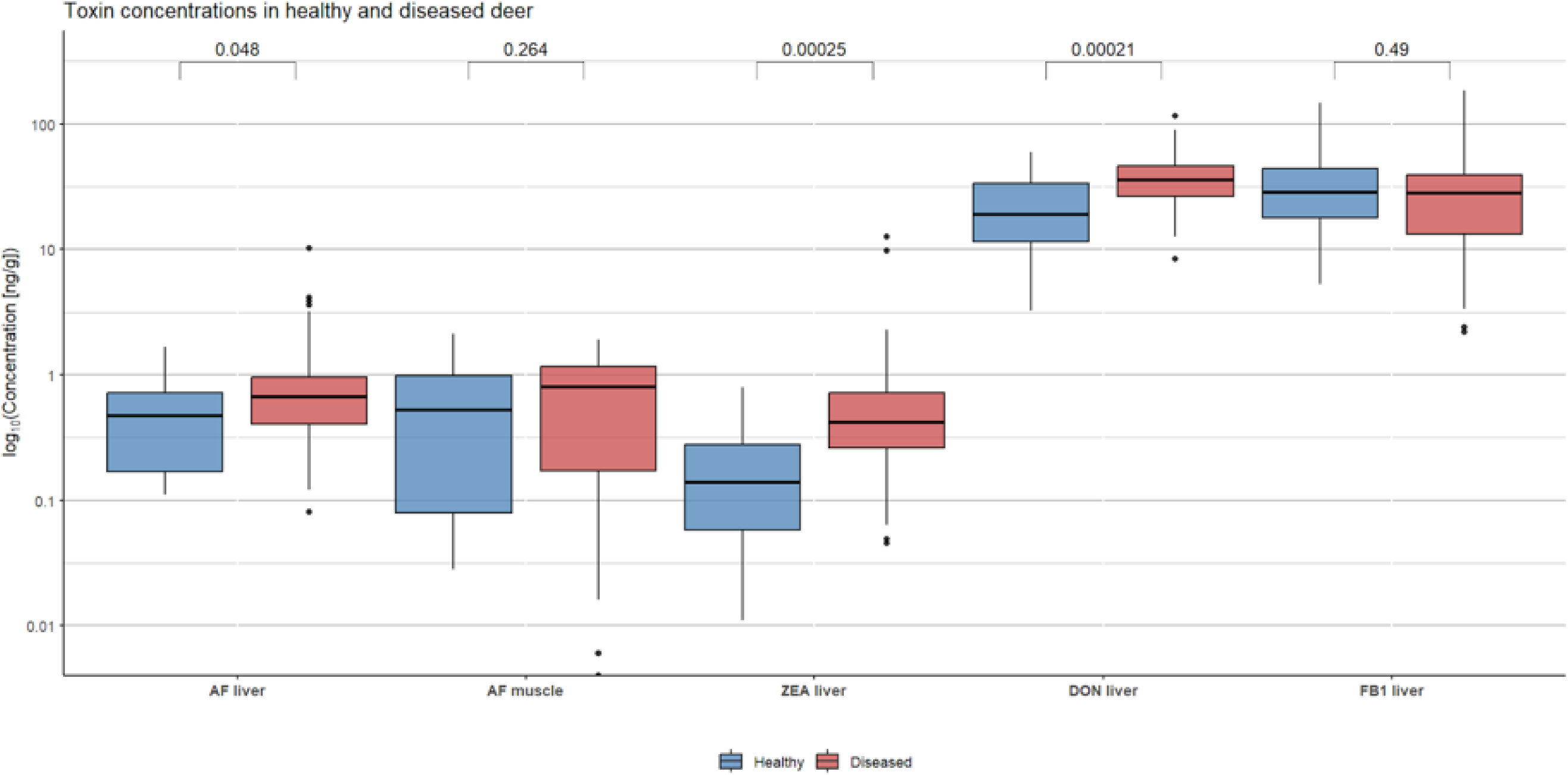

**Supplementary Data 10.**
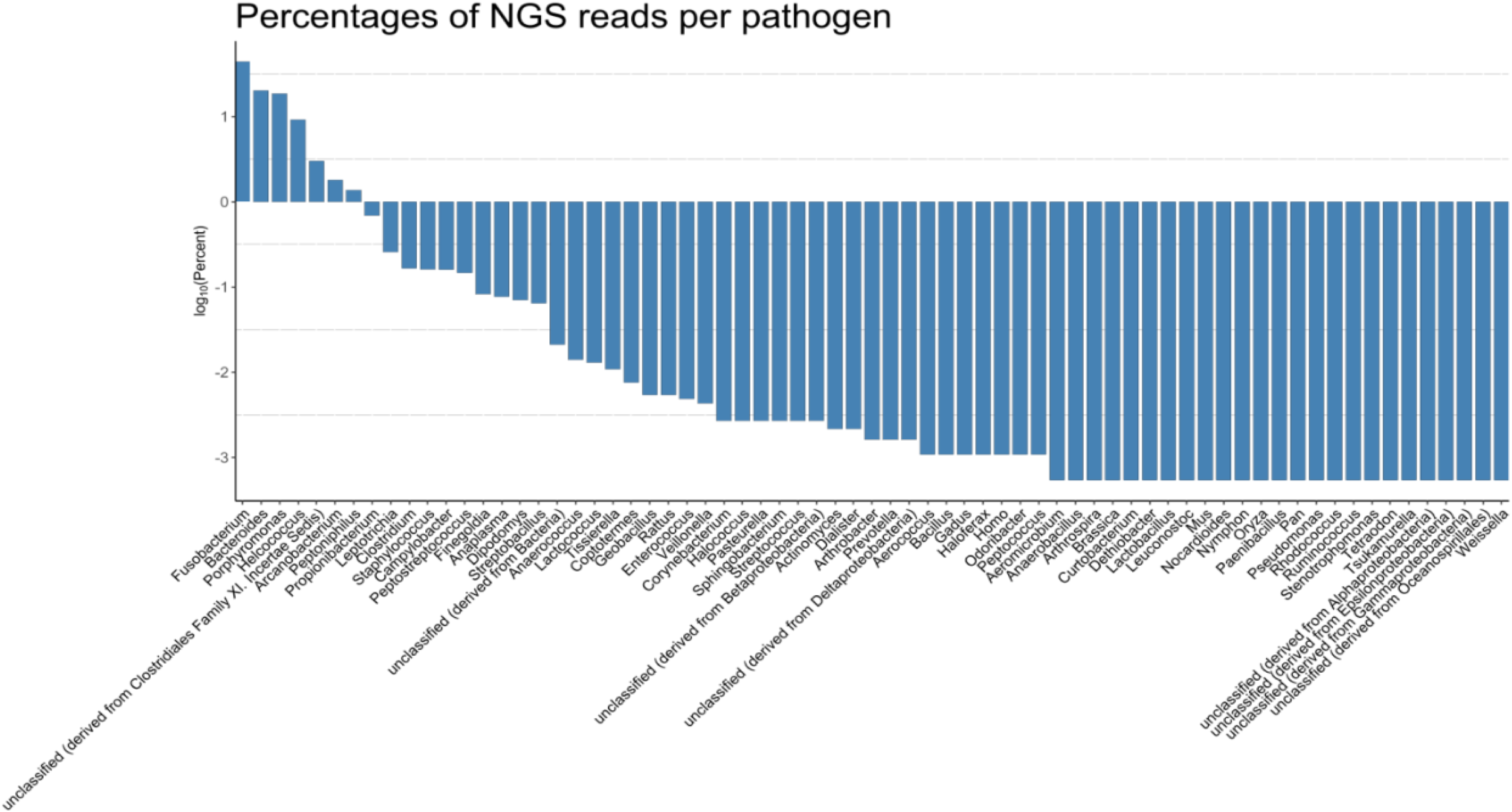

